# DNA methylation by CcrM contributes to genome maintenance in the *Agrobacterium tumefaciens* plant pathogen

**DOI:** 10.1101/2023.11.28.569018

**Authors:** Sandra Martin, Florian Fournes, Giovanna Ambrosini, Christian Iseli, Karolina Bojkowska, Julien Marquis, Nicolas Guex, Justine Collier

## Abstract

The <u>c</u>ell <u>c</u>ycle-regulated DNA <u>m</u>ethyltransferase CcrM is conserved in most *Alphaproteobacteria*, but its role in bacteria with complex or multicentric genomes remains unexplored. Here, we compare the methylome, the transcriptome and the phenotypes of wild-type and CcrM-depleted *Agrobacterium tumefaciens* cells with a dicentric genome with two essential replication origins. We find that DNA methylation has a pleiotropic impact on motility, biofilm formation and viability. Remarkably, CcrM promotes the expression of the *repABC^Ch2^* operon, encoding proteins required for replication initiation/partitioning at *ori2*, and inhibits *gcrA*, encoding a conserved <u>g</u>lobal <u>c</u>ell cycle regulator. Imaging *ori1* and *ori2* in live cells, we show that replication from *ori2* is often delayed in cells with a hypo-methylated genome, while *ori2* over-initiates in cells with a hyper-methylated genome. We thus propose that methylation by CcrM stimulates RepABC-dependent chromosomal origins, uncovering a novel and original connection between CcrM-dependent DNA methylation and genome maintenance in an *Alphaproteobacterial* pathogen.

## INTRODUCTION

Approximately 10% of the sequenced bacterial strains display complex genomes with more than one essential replicon and/or more than one essential replication origin. These include many *Alphaproteobacteria* that act as human, animal or plant pathogens, such as *Brucella abortus* or *Agrobacterium tumefaciens,* and that have very diverse modes of life ^1,2^. Multipartite or multicentric genomes pose significant challenges for genome stability and survival over generations, as the replication and the partitioning of multiple essential origins need to be coordinated with one another and with other events of the cell cycle. Although *Alphaproteobacteria* represent ∼26% of the sequenced bacterial strains with multipartite genomes ^3^, mechanisms they use to ensure the maintenance of multipartite/multicentric genomes are however largely underexplored compared to *Gammaproteobacteria ^4–6^*. In *Alphaproteobacteria*, replication of the main chromosome is supposedly always dependent on the very conserved initiator of DNA replication DnaA that binds to multiple sites on relatively classical bacterial origins of replication ^7^. Instead, replication of secondary chromosomes (carrying essential genes) or mega-plasmids is usually dependent on *repABC* operons ^4,5^. In such cases, RepC acts as the initiator of DNA replication by binding to an origin that is usually directly located inside the *repC* gene ^4,8^. Then, RepA and RepB (homologs of ParA and ParB) contribute to replicon partitioning through their binding to *parS* sequences located inside and/or next to the cognate *repABC* operon ^9^. Interestingly, *repABC* loci usually carry a higher-than-expected number of 5’-GANTC-3’ motifs that are often located in the promoter region driving the transcription of *repABC* operons, in putative origins located inside *repC* genes and in putative *repE* promoter regions driving the transcription of RepE small regulatory RNAs that can down-regulate *repC* transcription and translation ^4,10^. In nearly all *Alphaproteobacteria* except *Rickettsiales* and *Magnetococcales*, adenines located in such GANTC motifs are methylated (m6A) by the <u>c</u>ell <u>c</u>ycle-regulated DNA <u>m</u>ethyltransferase (MTase) CcrM ^11^. This solitary MTase, which is not associated with a cognate endonuclease, was initially discovered in the *Caulobacter crescentus Alphaproteobacterium* that has a single circular chromosome. In this bacterium, CcrM was shown to be present and active only at the very end of the S-phase of the cell cycle in pre-divisional cells ^12,13^. As a consequence, newly replicated GANTC motifs spread throughout the genome stay hemi-methylated for a significant period of the cell cycle after the passage of the replication fork, especially if these motifs are located closer to the origin of replication than to the terminus of replication of its unique chromosome ^14^. In *C. crescentus*, the methylation of hundreds of promoter regions by CcrM has a major impact on its transcriptome ^11^, notably because the co-conserved <u>g</u>lobal <u>c</u>ell cycle regulator GcrA can sense such epigenetic signals to modulate gene expression ^15–18^. One such gene strongly activated through methylation is the *ftsZ* gene required for cell division ^11,19^. Molecular genetics and experimental evolution analyses demonstrated that the essentiality of *ccrM* in fast-growing *C. crescentus* is specifically dependent on this key epigenetic activation of *ftsZ* expression ^19,20^. In *Brevundimonas subvibrioides*, where *ccrM* is not essential, the CcrM and GcrA regulons are significantly different compared to those of *C. crescentus ^21^*, demonstrating a relatively fast evolution of epigenetic regulatory pathways even in closely related *Alphaproteobacteria*. However, so far, studies did not focus on the impact of CcrM-dependent methylation in *Alphaproteobacteria* with more complex or multipartite genomes, despite observations suggesting that CcrM is probably also essential in *B. abortus ^22^* and *A. tumefaciens ^23,24^*. In the study described here, we aimed at filling this gap using the *A. tumefaciens* plant pathogen, which is the causal agent of the crown gall disease and a live biotechnological tool used for the genetic manipulation of plants ^25^. Earlier findings indicated that CcrM is also a cell cycle-regulated DNA MTase in this *Alphaproteobacterium ^23^*. *A. tumefaciens* displays a complex genome with two essential (*ori1* and *ori2*) and two dispensible (*ori^pAt^* and *ori^pTi^*) origins ^24,26,27^. In the original C58 strain sequenced in 2001, it was shown that its genome consists in one circular chromosome (Ch1 with a DnaA-dependent *ori1*), one linear chromosome (Ch2 with a RepC^Ch2^-dependent *ori2*) and two mega-plasmids encoding important virulence factors (pTi and pAt, with RepC^pTi^/RepC^pAt^-dependent *ori^pTi^* and *ori^pAt^*, respectively) ^26,27^. However, a very recent study from 2022 showed that many of the C58 strains used in laboratories over the world carry a unique dicentric linear chromosome instead of two distinct chromosomes (Fig.1a, left side) ^28^. Still, even in such strains, the *repABC^Ch2^* module (including *ori2)* appears to remain essential for survival since it could not be disrupted during Tn-seq experiments and since the *repB^Ch2^* gene could not be deleted ^28^, showing that initiation at *ori2* and/or *ori2* partitioning are essential in the two described *A. tumefaciens* C58 strains.

**Figure 1:**
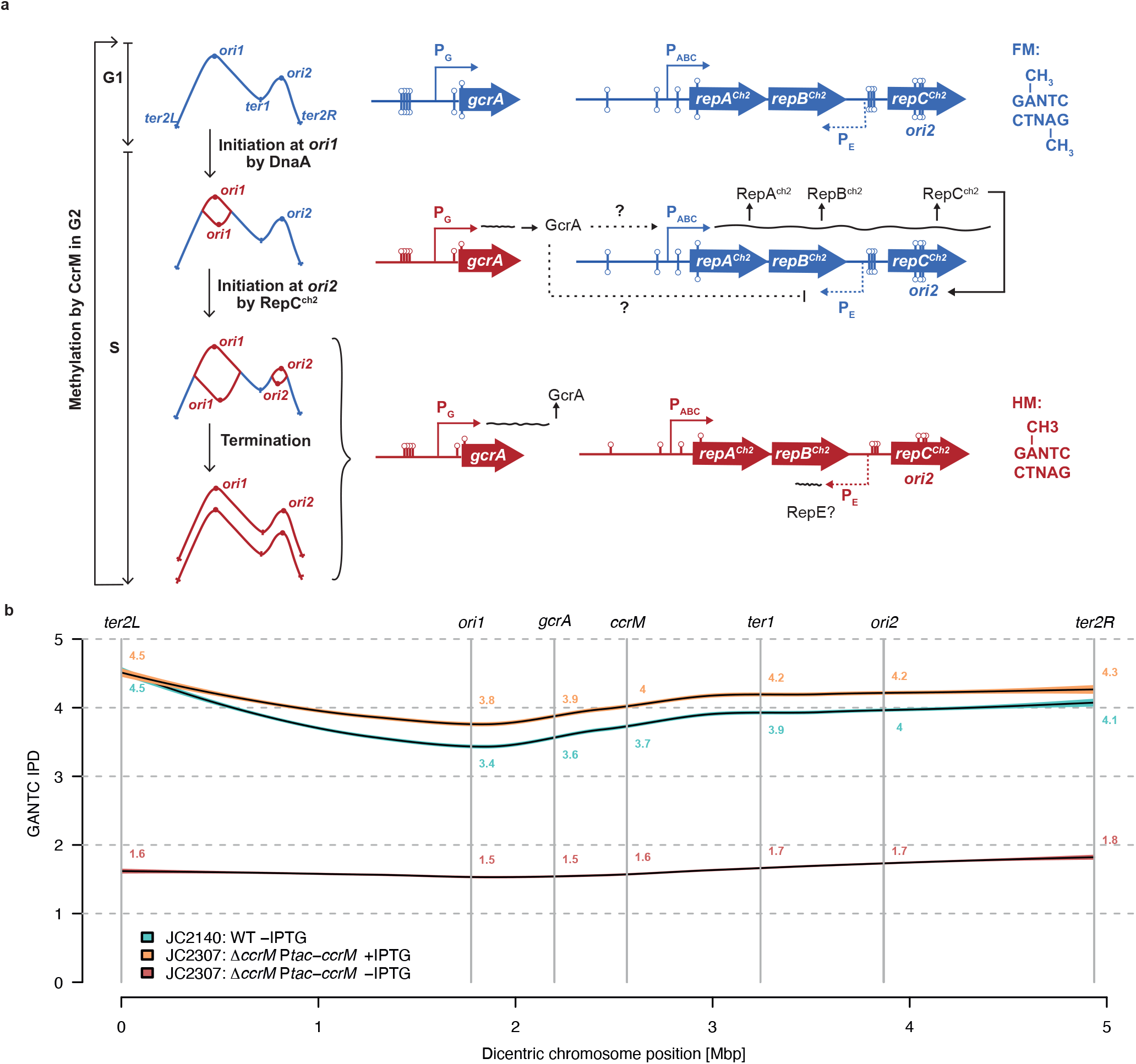
Methylation of GANTC motifs on the dicentric chromosome of *A. tumefaciens*. **(a)** Schematic showing the predicted methylation state of GANTC motifs depending on their chromosomal location and on cell cycle progression. “FM” stands for fully-<u>m</u>ethylated DNA (blue color) with m6A on both GANTC strands and “HM” stands for <u>h</u>emi-<u>m</u>ethylated DNA (red color) with m6A on only one GANTC strand (the newly-replicated strand is not yet methylated by the CcrM MTase). The right side of this schematic includes a working model concerning the impact of *gcrA* and *repABC^Ch2^* loci methylation by CcrM on their expression/activities as a function of cell cycle progression based on new findings from this study. Lollipops indicate m6A in GANTC motifs (including 200 bp upstream of ORFs). Wavy lines indicate when transcription from P_G_ (promoter upstream of *gcrA*), P_ABC_ (promoter region upstream of *repABC^Ch2^*) or P_E_ (putative promoter driving RepE expression) is expected to take place based on their methylation states. **(b)** CcrM-dependent methylation of GANTC motifs on the *A. tumefaciens* dicentric chromosome. gDNA samples from WT and JC2307 (Δ*ccrM* P*tac-ccrM*) cells that grew exponentially in ATGN+/-IPTG for 7 hours were analyzed by SMRT-seq. GANTC IPD ratios were directly generated by the PacBio software. A smooth curve was then fitted between these ratios and the dicentric chromosome position using the LOESS function of R with default parameters. The 95% confidence interval was then plotted to predict average GANTC IPD ratios depending on their position on the dicentric chromosome from *ter2L* (position “0 Mbp”) to *ter2R* (position “4.936 Mbp”).

Here, we constructed a conditional *ccrM* mutant and combined methylome and transcriptome analyses, together with single cell reporter assays, to test the impact of CcrM-dependent methylation on the expression and the maintenance of the *A. tumefaciens* complex genome. Our results demonstrate that CcrM-dependent methylation is an essential process for fast-growing *A. tumefaciens* cells and that it has a major impact on the global transcriptome including on the expression of its three *repABC* operons. Consistent with an epigenetic control of chromosome replication/partitioning in such *Alphaproteobacteria*, we also found that the timing or the frequency of *ori2* firing/partitioning are affected when the *A. tumefaciens* genome becomes hypo-or hyper-methylated due to genetic perturbations. Altogether, we thus uncovered an original connection between CcrM-dependent methylation and genome maintenance in a pathogenic *Alphaproteobacterium* with an essential RepABC-dependent replicon.

## RESULTS

### CcrM is essential in *A. tumefaciens* cells cultivated in complex media

Considering the predicted essentiality of *ccrM* in *A. tumefacien ^23,24^*, we engineered a conditional *ccrM* mutant to be able to analyze the methylome, the transcriptome and the phenotypes of cells as CcrM gets depleted. This enabled us to obtain information on the impact of DNA methylation in *A. tumefaciens*. In practice, a second copy of *ccrM* (*Atu0794*) under the control of the IPTG-inducible P*tac* promoter was introduced into the dicentric chromosome of an *A. tumefaciens* C58 derivative strain, and then the native *ccrM* gene was deleted following a standard double recombination procedure ^29^, generating strain JC2307 *(ΔccrM* P*tac-ccrM*) on ATGN minimal medium containing IPTG. Even if colonies grew very slowly on ATGN lacking IPTG compared to ATGN with IPTG (Fig.2a), colonies were still largely detectable, indicating either that CcrM was not efficiently depleted in the absence of the IPTG inducer, or that *ccrM* is not strictly essential under such growth conditions. Consistent with this first observation, we also found that JC2307 cells could grow relatively well in liquid ATGN medium, even if their growth rate progressively decreased over time after the removal of the IPTG inducer (Fig.2b). Colony forming unit (CFU) assays also showed that cell viability remained normal for extended periods of time (Fig.S1), even if CcrM levels already became very hard to detect from cell extracts using immunoblotting experiments after 7 hours of growth in ATGN without IPTG (Fig.S2). The morphology of CcrM-depleted cells (cultivated in ATGN without IPTG) appeared as relatively similar to control cells (Fig.2c), even if quantitative analyses showed that such cells were slightly shorter than control cells expressing *ccrM* (median length of 1.64 μm compared to 1.94-1.99 μm) (Fig.S3, ATGN).

Strikingly, the impact of CcrM depletion was much stronger in cells cultivated in complex media. Indeed, JC2307 cells could hardly form colonies on YEB or LB plates lacking IPTG (Fig. 2a), had severe difficulties to grow in liquid YEB or LB media without IPTG (Fig.2b) and appeared as significantly elongated and/or enlarged (Fig.2c and Fig.S3) before they started to lyse/die (Fig.S1). Time-lapse microscopy experiments also showed that most JC2307 cells cultivated onto YEB agarose pads lacking IPTG lysed or developed abnormal morphologies (swollen, elongated and multi-polar cells) over time (Fig.S4 and Movies S1&S2). Importantly, the presence of IPTG in these media to induce P*tac-ccrM* led to a complete (in YEB) or partial (in LB) complementation of these phenotypes (Fig.2 and Fig.S1&S3), showing that they were, indeed, connected to insufficient levels of CcrM and not to polar effects.

**Figure 2:**
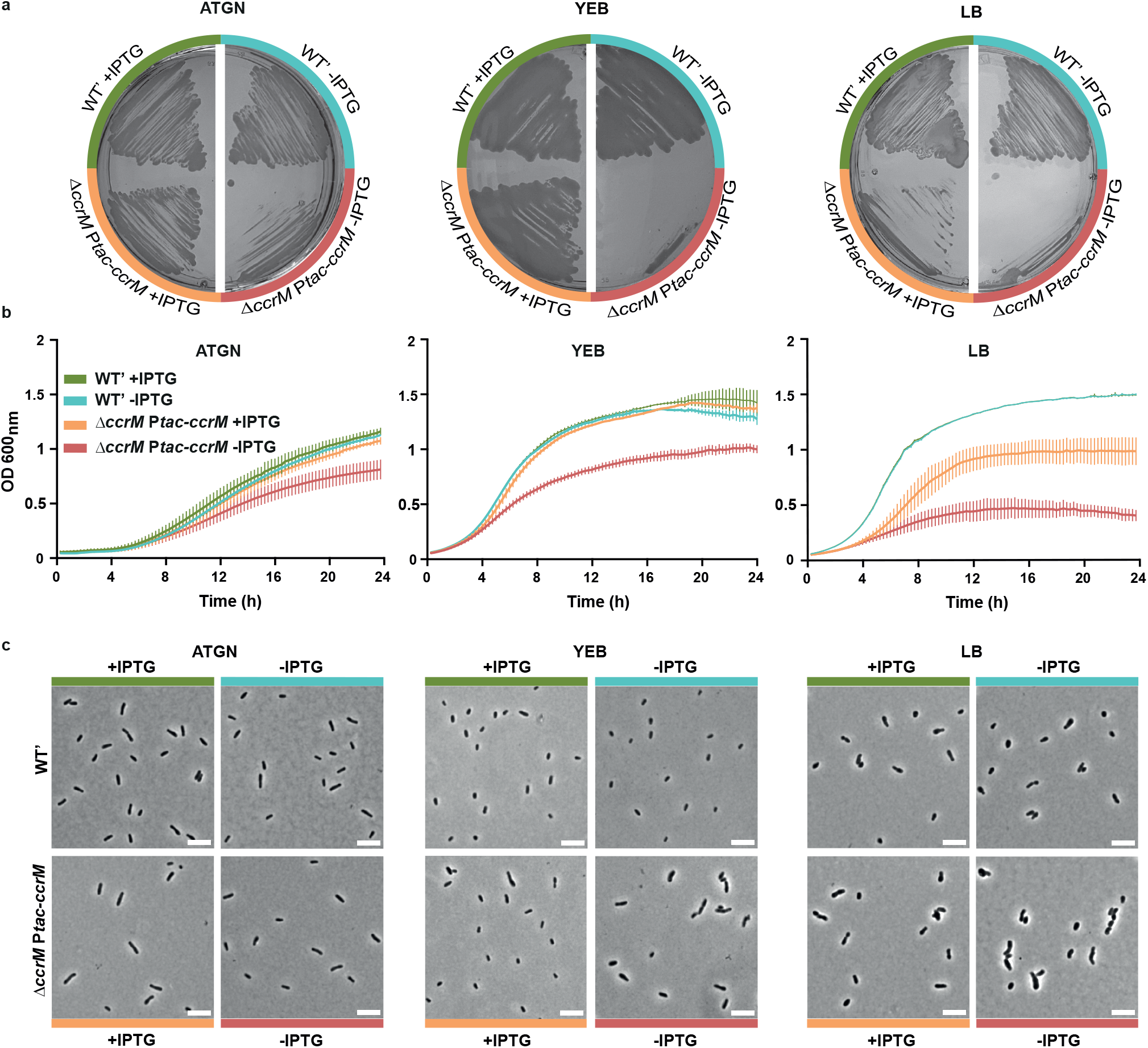
Depletion of CcrM results in slow growing, elongated and swollen cells when cultivated in complex media. **(a)** JC2141 (WT’) and JC2307 (Δ*ccrM* P*tac-ccrM*) cells were plated onto ATGN, YEB or LB agarose plates containing or not the IPTG inducer and colonies were then cultivated for ∼3 days prior to imaging. **(b)** JC2141 and JC2307 strains were pre-cultured into liquid ATGN, YEB or LB media + IPTG. Cells from over-night cultures were then washed and diluted to an OD_600_∼0.1 into the indicated medium and placed into a plate reader to record the OD_600_ every 15 minutes. Error bars represent means +/- standard deviations from three biologically independent replicates. Note that the two WT’ curves overlap for the LB growth condition. **(c)** JC2141 and JC2307 strains were pre-cultured into liquid ATGN+IPTG and then diluted into ATGN or YEB or LB with IPTG. Once cultures reached exponential phase again, cultures were washed and cells were resuspended to an OD_600_∼0.2 into ATGN, YEB or LB +/-IPTG at time T0. Cells were then fixed and imaged by phase-contrast microscopy after 24h of growth. White scale bars inside images are 5μm.

Altogether, the use of this conditional mutant for phenotypic analyses demonstrated that CcrM is essential for the survival of *A. tumefaciens* cells cultivated in or on complex media, but that it may be dispensible, or at least not required at similar levels, in cells cultivated in or on minimal media.

### CcrM-depleted cells cultivated in minimal medium have a hypo-methylated genome and display motility and adhesion defects

To test if CcrM levels become too limiting to ensure an efficient methylation of genomic GANTC motifs in the conditional mutant cultivated in minimal medium (when cells remain viable), we performed two assays. First, we compared the efficiency of digestion by HinfI of gDNA samples prepared from WT or JC2307 cells cultivated in ATGN with or without IPTG. HinfI is a methylation-dependent endonuclease that can only cut non-methylated GANTC motifs ^23^. Consistent with the known efficient activity of CcrM in *A. tumefaciens*, we confirmed that restriction of the WT genome by HinfI was essentially undetectable (Fig.S5). In contrast, the genome of JC2307 cells cultivated in ATGN without IPTG for 7 hours became highly sensitive to HinfI digestion (Fig.S5), indicating that a majority of its double-stranded GANTC motifs become un-methylated when CcrM is depleted. To get a more quantitative evaluation of the methylation state of the *A.tumefaciens* genome upon CcrM depletion, we next analyzed the methylome of cells cultivated in these same growth conditions using single molecule real-time sequencing (SMRT-Seq), a method that is now commonly used to distinguish methylated (m6A) from non-methylated adenines on bacterial genomes based on measures of interpulse duration (IPD) ^30,31^. Using gDNA extracted from WT cells, we found that the average IPD ratio of adenines located in GANTC motifs located near *ori1* (IPD∼3.4) was lower than that of GANTC motifs located next to *ter2L* (IPD∼4.5) on the dicentric chromosome (Fig.1b), which is fully consistent with the known cell cycle regulation of CcrM-dependent methylation in *A. tumefaciens* cells (newly replicated double-stranded GANTC motifs staying hemi-methylated until the end of the S-phase of the cell cycle; Fig.1a)^23^. In contrast, using gDNA from JC2307 (*ΔccrM* P*tac-ccrM*) cells cultivated in ATGN without IPTG for 7 hours, the average IPD ratio of adenines located in GANTC motifs throughout the dicentric chromosome dropped to an average value of ∼1.6 (compared to an average of ∼3.8 for WT cells and ∼4.1 for *ΔccrM* P*tac-ccrM* cells cultivated in ATGN with IPTG) (Fig.1b and Fig.S6). This observation confirmed that a vast majority of the 5166 double-stranded GANTC motifs found on the dicentric chromosome of JC2307 cells are in a non-methylated state after 7 hours of CcrM depletion. Importantly, finding viable conditions (Fig.2 & Fig.S1) when *A. tumefaciens* cells display a largely hypo-methylated genome (Fig.1b & Figs.S5&S6) made it possible to test the impact of DNA methylation by CcrM on the transcriptome of *A. tumefaciens* cells with a complex multicentric genome.

Interestingly, even if such mutant cells did not display obvious viability, growth or morphology defects in ATGN medium (Fig.2 and Figs. S1&S3), we still noticed two interesting phenotypes. First, cells displayed a significant swarming defect as could be seen with the ∼35% decrease in swarming area when placed onto semi-solid ATGN medium without IPTG compared to semi-solid ATGN with IPTG (Fig.3a). Second, we observed a ∼35% decrease in adhesion/biofilm formation in ATGN without IPTG (Fig. 3b) as could be measured using static coverslip assays with crystal violet ^29^.

**Figure 3:**
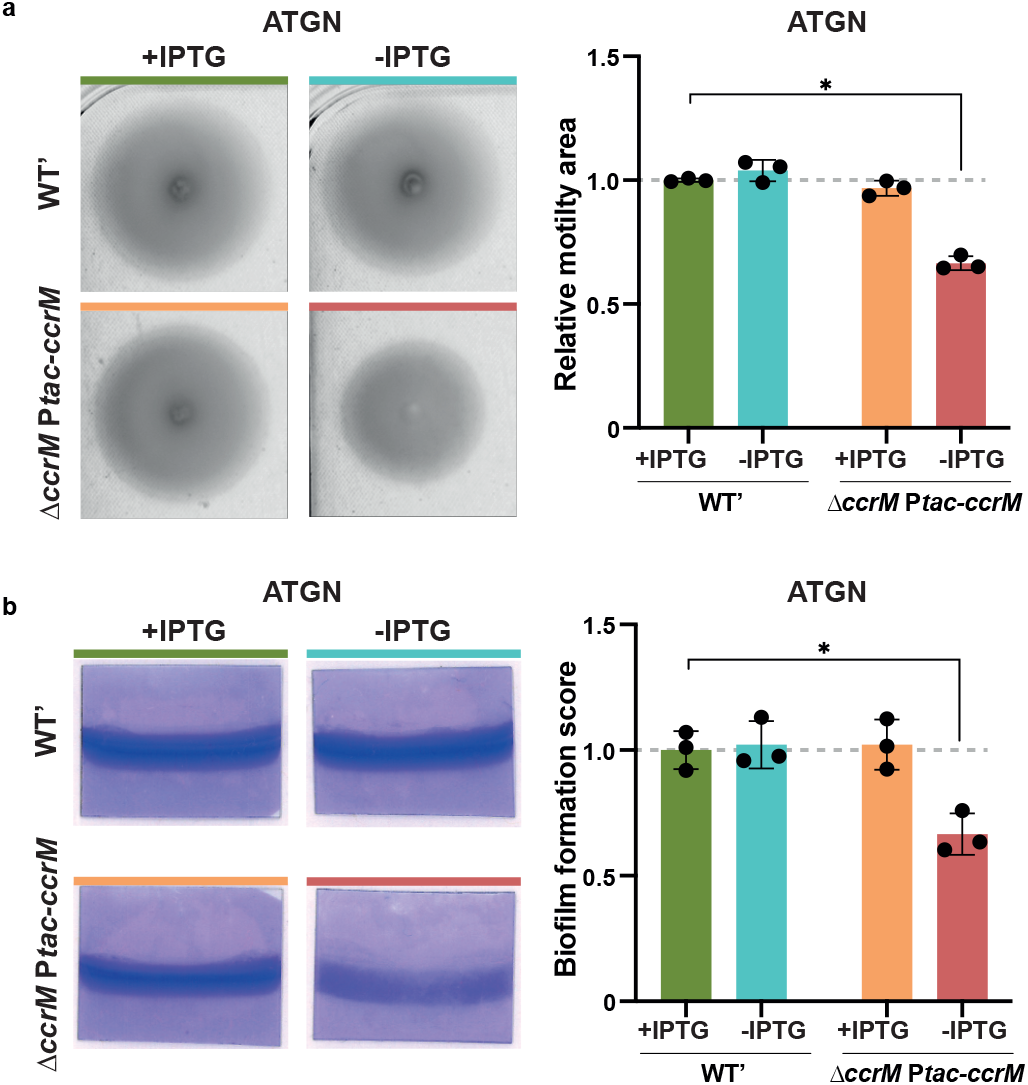
Depletion of CcrM results in motility and biofilm formation defects in minimal medium. **(a)** CcrM-depleted cells are less motile on soft-agar plates. JC2141 (WT’) and JC2307 (Δ*ccrM* P*tac-ccrM*) strains were pre-cultured into liquid ATGN+IPTG. Over-night cultures were then washed and adjusted to an OD_600_∼0.6 into ATGN. 5μL were then spotted onto ATGN soft-agar plates +/-IPTG and plates were incubated for 48 hours prior to imaging. Representative images are shown on the left. Quantitative analysis using images from 3 biological replicates with 2 technical replicates each are shown on the right (relative to WT’ cells cultivated in ATGN+IPTG). Error bars represent means +/- standard deviations from the three biologically independent replicates (shown as black circles). The only significant difference (unpaired student’s t-test) compared to WT’ cells cultivated into ATGN+IPTG soft-agar is indicated by * (P<0.01). **(b)** CcrM-depleted cells display attachment/biofilm formation defects. JC2141 and JC2307 strains were pre-cultured into liquid ATGN+IPTG. Over-night cultures were then washed and adjusted to an OD_600_∼0.05 into ATGN+/-IPTG. Cultures were then assayed for biofilm formation on vertical plastic coverslips immersed into the culture. Representative images of coverslips after a 72-hour incubation period at room temperature are shown on the left. Biofilm scores were then measured using data from 3 biological replicates and are shown on the right (relative to WT’ cells cultivated in ATGN+IPTG). Error bars represent means +/- standard deviations from the three replicates (shown as black circles). The only significant difference (unpaired T-test) compared to WT’ cells cultivated into ATGN+IPTG soft-agar is indicated by * (P<0.01).

Altogether, these methylome and phenotypic assays showed that *A. tumefaciens* cells with hypo-methylated GANTC motifs on their genome are viable in minimal medium, but still display detectable phenotypes suggesting that some genes may be mis-expressed as a result of their hypo-methylation.

### DNA methylation by CcrM has a major impact on the *A. tumefaciens* transcriptome

To collect information on the origin of the observed phenotypes (Fig.2&3) and gather potential cues on why DNA methylation by CcrM becomes essential in cells cultivated in complex media (Fig.2 and Fig.S1), we next decided to compare the transcriptome of viable *A. tumefaciens* JC2307 cells soon after the drop in detection of methylated GANTC motifs (7 hours in ATGN without IPTG) with that of mutant cells cultivated in ATGN with IPTG or WT cells (Fig.1b). RNA-Seq experiments revealed a significant (adjusted P-value <0.01) and strong (fold change > 2) impact on the expression of 273 genes (Fig.4a and Table S4): 59 genes were down-regulated, while 214 genes were up-regulated in response to CcrM depletion (comparing JC2307 cells cultivated with or without IPTG for 7 hours). Instead, the transcriptome of CcrM-repleted cells (JC2307 cells cultivated with IPTG), appeared as extremely similar to the transcriptome of WT cells (Fig.S7 and Table S4), showing that expression of *ccrM* from the P*tac* promoter instead of from its native promoter does not lead to significant gene mis-regulation. Among the 273 genes that were significantly mis-regulated upon CcrM depletion, 75 genes (27%) displayed at least one GANTC motif in their putative promoter region (200 bp upstream of each ORF), including 22 of the 59 (37%) down-regulated genes and 53 of the 214 (25%) up-regulated genes (Fig.4a and Table S5).

As a first point of interest, we compared this potential “direct regulon” of CcrM (75 genes) in *A. tumefaciens* (Table S5) with previously published “direct regulons” of CcrM in the distantly related *C. crescentus* (152 genes) and *B. subvibrioides* (129 genes) *Alphaproteobacteria*. This analysis showed that only six genes of the *A. tumefaciens* CcrM “direct regulon” had orthologs that also belonged to the *C. crescentus* or *B. subvibriodes* “direct regulons” (Table S5). Thus, we concluded that epigenetic mechanisms of regulation evolved tremendously between *Alphaproteobacteria* with different genome architectures and lifestyles.

Another striking new finding was that the *A. tumefaciens* CcrM regulon included six genes belonging to the three *repABC* operons (essential for the replication of the dicentric chromosome and of the two mega-plasmids) among the genes that were significantly down-regulated in CcrM depleted cells (Fig.4 and Table S5). In addition, many genes encoding proteins potentially involved in SOS-related responses to DNA damage or replication defects (examples: RecA, RecQ, three orthologs of DNA Pol Y and three orthologs of ImuA) were significantly up-regulated. Noteworthy, the *gcrA* homolog of *A. tumefaciens (Atu0426),* encoding a putative methylation-sensitive global regulator, was strongly activated (2.8-fold induction) upon CcrM depletion, while the *ftsZ^At^* (*Atu2086/ftsZ2*) gene required for *A. tumefaciens* cell division remained un-affected (Fig.4 and Table S4) confirming strong differences with what was previously observed in *C. crescentus ΔccrM* mutant cells ^11,19^. The impact of CcrM depletion on the expression of this interesting selection of genes was also verified by qRT-PCR (Fig.4b) using RNA samples prepared not only from mutant cells cultivated in ATGN +/- IPTG for 7 hours (same conditions as the RNA-Seq experiments), but also from mutant cells cultivated in YEB +/- IPTG for 5.5 hours (prior to the detection of cell death without the IPTG inducer of P*tac-ccrM* as shown in Fig.S1). These experiments showed that DNA methylation by CcrM also promotes *repABC^Ch2^* expression and represses *gcrA* expression in cells cultivated in complex YEB medium, while it does not have a significant impact on the expression of the essential *ftsZ^At^* gene under such growth conditions (Fig.4b, right panel).

**Figure 4:**
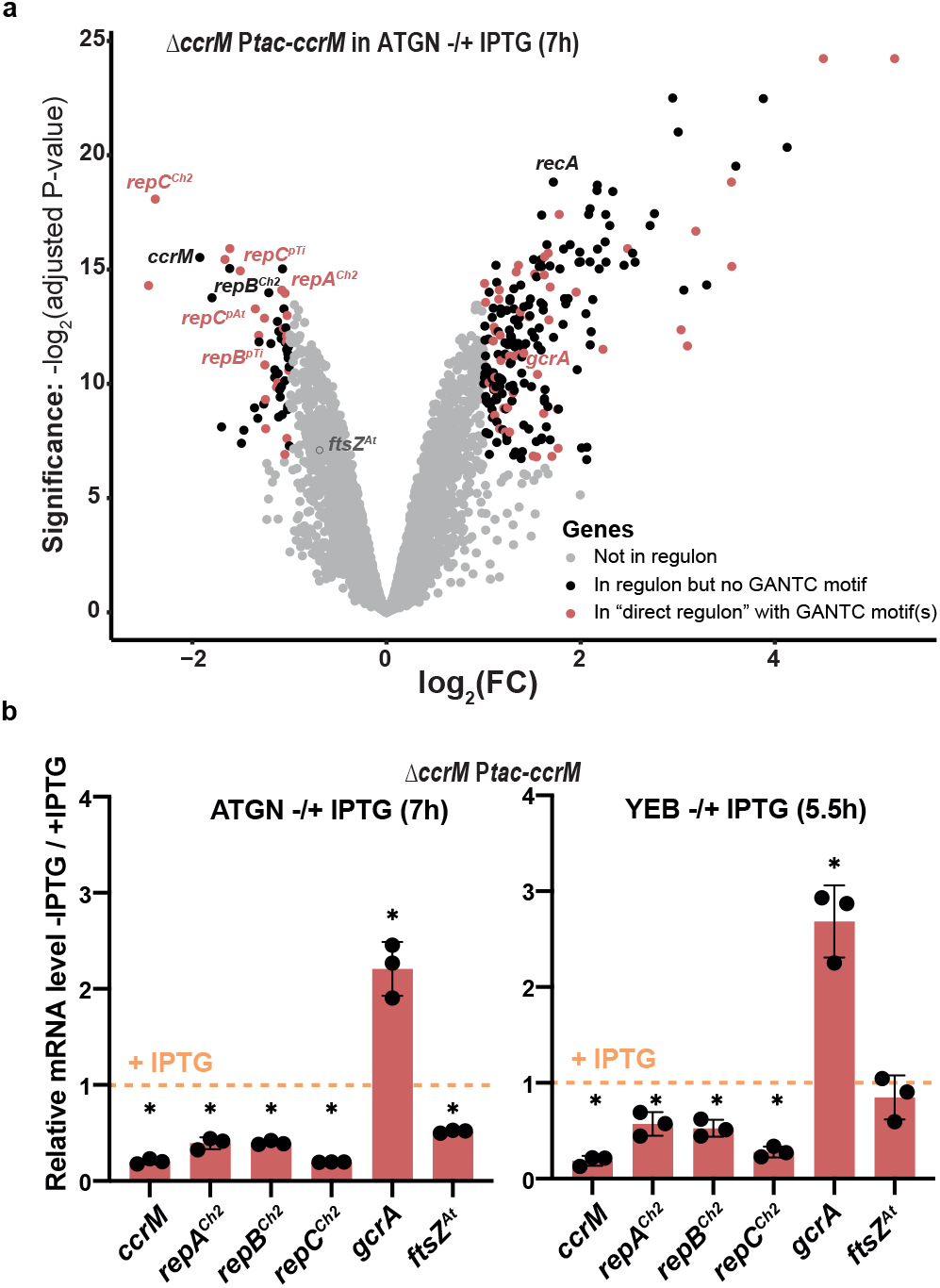
Depletion of CcrM has a major impact on the *A. tumefaciens* transcriptome in cells cultivated for 7 hours in minimal medium. **(a)** Volcano plot showing RNA-Seq results and analyses comparing the transcriptome of exponentially growing JC2307 (Δ*ccrM* P*tac-ccrM)* cells cultivated in ATGN+/-IPTG for 7 hours (same cultures as those used for methylome experiments described in Fig.1b). FC indicates the fold-change when comparing the -IPTG condition with the +IPTG condition. Each dot corresponds to a gene. Grey dots correspond to genes that are not considered as differentially regulated (= FC<2 or adjusted P-value >0.01). Black dots correspond to 198 genes that are differentially regulated (FC> 2 and adjusted P-value <0.01) but that do not contain a GANTC motif in the 200 bp region upstream of the ORF. Red dots correspond to the 75 genes that are differentially regulated (FC> 2 and adjusted P-value <0.01) and that do contain minimum one GANTC motif in the 200 bp region upstream of the ORF (“direct regulon”). The precise identity/annotation/COG category/GANTC motif locations for all genes can be found in Tables S4&S5. A Glimma Volcano Plot (interactive HTML graphic) is also available in the supplementary information. Adjusted P-values were calculated based on three independent biological replicates for each strain/growth condition. **(b)** qRT-PCR results confirming the impact of CcrM on the expression of a selection of genes in JC2307 (Δ*ccrM* P*tac-ccrM*) cells cultivated in minimal (left) and complex (right) media. The graphs show relative mRNA levels in cells cultivated without the IPTG inducer, compared to cells cultivated with the IPTG inducer (then set to an arbitrary value of 1 for each gene = orange dotted line). The same RNA samples as those used in (a) were used for the “ATGN+/-IPTG (7h)” panel. RNA samples used for the “YEB+/- IPTG (5.5h)” panel were prepared from JC2307 cells pre-cultured in ATGN+IPTG and then diluted into YEB+IPTG for growth until exponential phase. Cells were then washed and resuspended into YEB+/-IPTG at Time 0. RNA samples were prepared after 5.5h of growth. For both panels, three biological replicates with three technical replicates each were used. Significant differences (Wilcoxon rank sum test) when comparing -IPTG/+IPTG are indicated by * (P<0.0001).

These exciting discoveries led us to hypothesize that CcrM-depleted cells may express sub-optimal levels of the essential RepC^Ch2^ *ori2* initiator and RepAB^Ch2^ *ori2* partitioning proteins, leading to potential replication delays from *ori2* and the resulting onset of a mild SOS response in a subset of cells cultivated in ATGN medium. This may also relate to the essentiality of *ccrM* in fast-growing *A. tumefaciens* cells (in YEB and LB media) (Fig.2).

### Replication/partitioning of *ori2* is decoupled from replication/partitioning of *ori1* in CcrM-depleted cells

A few recent studies used live cell fluorescence microscopy experiments to visualize the number and the sub-cellular localization of *ori1* and *ori2* as a function of cell cycle progression (cell length). These studies showed that *ori1* and *ori2* colocalize at the old pole of newborn G1 phase cells ^32^ (Fig.5a). At the onset of the S-phase of the cell cycle, replication first starts at *ori1* and then at *ori2* following a significant delay. Once duplicated, one copy of each origin (*ori1* and then *ori2*) moves towards the new cell pole ^33^ (Fig.5a). Importantly, the timing of origin firing and the localization of origins are strikingly similar in C58 strains with two chromosomes compared with strains with one dicentric chromosome ^28^. We therefore decided to use these same fluorescent reporters to visualize the impact of CcrM depletion on the timing of *ori1* and *ori2* replication and on their partitioning during the *A. tumefaciens* cell cycle.

As a first test, we visualized *ori1* or *ori2* localization using a *ygfp-parB^pMT1^-parS^pMT1^* reporter integrated next to *ori1* or *ori2 ^33^*, respectively, in the dicentric chromosome of JC2307 (*ΔccrM* P*tac-ccrM*) cells. Fluorescence imaging of cells growing exponentially in ATGN +/-IPTG (Fig.S8a) and analyses of demographs (Fig.5b&c) revealed that the duplication/partitioning of *ori2* tends to take place in CcrM-depleted (ATGN-IPTG) cells that have reached a longer length compared to CcrM-repleted (ATGN+IPTG) cells (Fig.5c), while no obvious difference could be seen when looking at *ori1* duplication/partitioning (Fig.5b). Moreover, a lower proportion of CcrM-depleted cells (∼18%) displayed two *ori2* foci compared to CcrM-repleted cells (∼38%) (Fig.S8b), while CcrM depletion had an un-significant impact on the overall proportion of S-phase cells (∼39% instead of 43% of cells displayed 2 *ori1* foci) (Fig.S8a). These first observations suggested that *ori2* duplication or partitioning is/are specifically delayed in cells that display a hypo-methylated genome.

As a second test, we also co-visualized *ori1* and *ori2* in JC2307 cells using a first *ygfp-parB^pMT1^-parS^pMT1^* reporter near *ori1* and a second *mcherry-parB^P1^-parS^P1^* reporter near *ori2*^32^. Fluorescence imaging of cells growing exponentially in ATGN+/-IPTG (Fig.S9a) and analyses of demographs (Fig.S9b) confirmed that duplication/partitioning of *ori2* tends to take place in CcrM-depleted (ATGN-IPTG) cells that have reached a longer length compared to CcrM-repleted (ATGN+IPTG) cells. Quantitative analyses using this double-labelled strain further showed that the proportion of S-phase cells (with two *ori1* foci) also displaying two *ori2* foci dropped ∼2-fold (from ∼62% to ∼30%) when mutant cells were cultivated in ATGN-IPTG compared to ATGN+IPTG (Fig.5d), while the overall proportion of S-phase cells (with two *ori1* foci) remained essentially similar (from ∼58% to ∼52%) (Fig.S10). Altogether, these results indicate that replication initiation at *ori2* and/or *ori2* partitioning are/is specifically delayed in mutant *A. tumefaciens* cells displaying a hypo-methylated genome, coinciding with a lower expression of the *repABC^Ch2^* operon (Fig.4).

### Chromosome replication over-initiates from *ori2* in CcrM-overexpressing cells

The results described above suggest that DNA methylation by CcrM promotes DNA replication from *ori2* in *A. tumefaciens* cells. To confirm this hypothesis, we also looked at the impact of CcrM over-expression on the number of *ori1* and *ori2* foci per cell. We expect that CcrM over-expression from the pRX-P*tac-ccrM* vector leads to a significant hyper-methylation of the *A. tumefaciens* genome with most GANTC motifs being fully-methylated throughout the cell cycle as previously shown using related vectors ^23^. In ATGN medium containing IPTG, only ∼35% of the cells over-expressing CcrM displayed a single *ori2* focus, compared to ∼54% of the cells carrying the empty control vector, while differences concerning *ori1* appeared as non-significant (47% compared to 45% with P-value>0.05) (Fig.6a&b). This observation suggested that replication from *ori2* can start sooner after (or even before) replication from *ori1* in cells with a hyper-methylated genome compared to WT cells, which is supported by demographs comparing the timing of *ori1*/*ori2* duplication as a function of cell size/cell cycle progression (Fig.6c&d). Consistent with such a boost of replication from *ori2*, we also observed that ∼20% of the CcrM-overexpressing cells displayed more than two *ori2* foci per cell, something that only very rarely (∼2% of cells) happened in WT cells (Fig.6a&b). Thus, *A.tumefaciens* cells with a hyper-methylated genome regularly over-initiate DNA replication from *ori2*.

## DISCUSSION

In this study, we discovered that the CcrM DNA MTase, which is conserved in all known *Alphaproteobacteria* with multipartite genomes ^11^, plays a role in controlling the timing of *ori2* replication/partitioning and the frequency at which *ori2* is replicated in *A. tumefaciens* cells. Our observations indicate that *ori2* initiates/segregates with a significant delay during the *A. tumefaciens* cell cycle (Fig.5 and Fig.S8&S9) when its dicentric chromosome is hypo-methylated due to a depletion of CcrM (Fig.1b and Fig.S5&6). Conversely, *ori2* appears to over-initiate when CcrM is over-expressed, leading to cells with unusually high *ori2* copy numbers (Fig.6). Thus, we propose that the hemi-methylation of the *repABC^Ch2^/ori2* locus after the onset of replication from *ori2* may be involved in preventing new *ori2* firing before the very end of the S-phase of the cell cycle when CcrM re-methylates the whole genome (Fig.1a). This would be somewhat reminiscent of the control of DnaA-dependent origins by the Dam/SeqA couple in *Gammaproteobacteria ^34^*. Moreover, the impact of CcrM-dependent methylation on *ori2* control may also provide an explanation for the apparent essentiality of *ccrM* in cells cultivated in complex media (Fig.2 and Fig.S1) since initiation at *ori2* is essential in *A. tumefaciens ^28^*. Why this process becomes more critical under such growth conditions may, for example, relate to higher coordination requirements during fast growth or to shorter half-lives of critical regulators (CcrM, RepABC^Ch2^, …) leading to more critical depletions of such elements. Interestingly, such CcrM-dependent control of secondary chromosomal origins may be conserved far beyond *A. tumefaciens*, as a former study showed that CcrM over-expression leads to abnormally high intra-cellular DNA contents in the *B. abortus* human pathogen ^22^.

**Figure 5:**
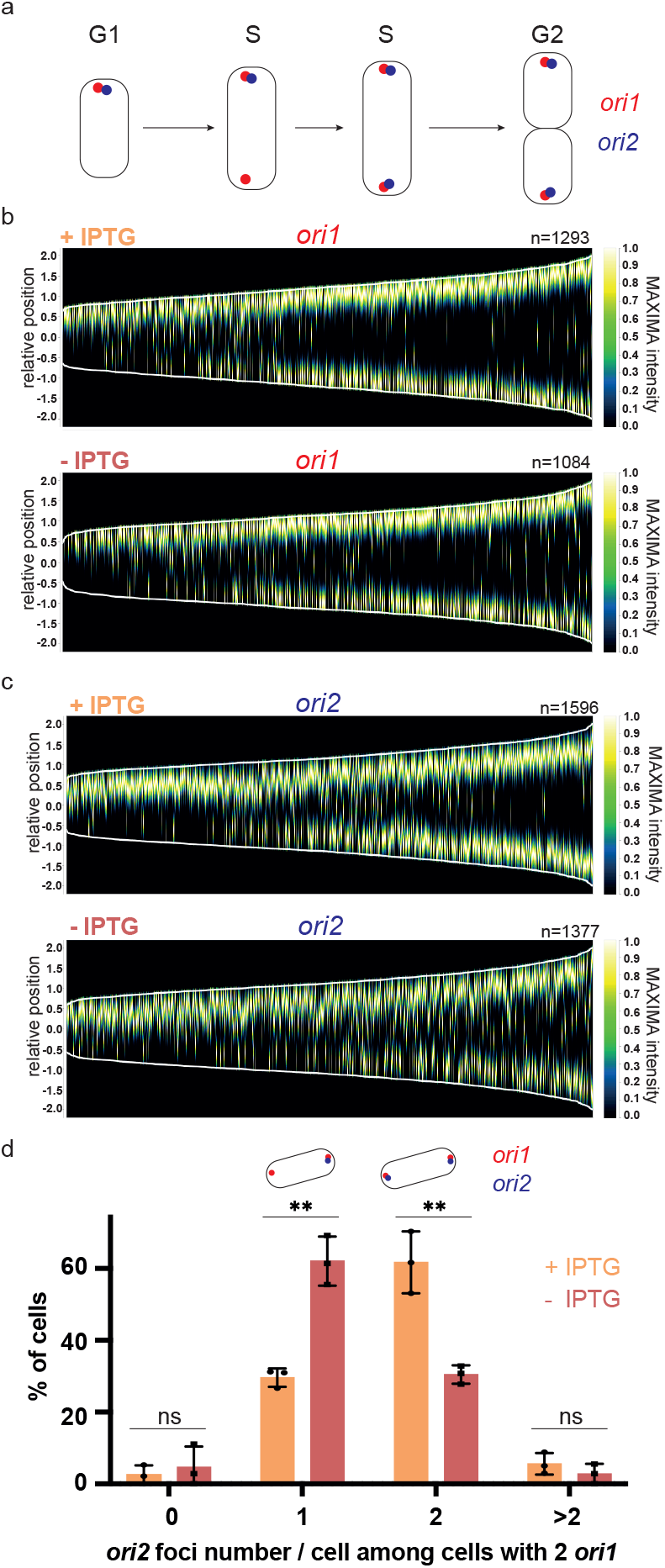
Depletion of CcrM results in *ori2* replication/partitioning delays in cells cultivated in minimal medium for prolonged periods of time. **(a)** Schematic representing the subcellular localization of *ori1* and *ori2* during the *A. tumefaciens* cell cycle. **(b)** and **(c):** Demographs showing the subcellular localization of *ori1* (b) or *ori2* (c) foci as a function of cell size (from images as shown in Fig.S8). JC2660 (*ΔccrM* P*tac-ccrM* with *ori1/ygfp* reporter in **(b)**) or JC2661 (*ΔccrM* P*tac-ccrM* with *ori2/ygfp* reporter in **(c)**) cells were cultivated over-night in ATGN+/-IPTG. Cultures were then diluted into ATGN+/-IPTG and grown exponentially for ∼6.5 hours (>15 hours into ATGN+/-IPTG). Relative position = 0 corresponds to mid-cell; only cells measuring from 1 to 4 μm-long were included into these demographs; n = number of cells used to construct each demograph. **(d)** Quantification of *ori2* number per cell among cells that are in S-phase (with two *ori1*). JC2836 (*ΔccrM* P*tac-ccrM* with *ori1/mcherry* and *ori2/ygfp* reporters) cells were cultivated over-night in ATGN+/- IPTG. Cultures were then diluted into ATGN+/-IPTG and grown exponentially for ∼6.5 hours (>15 hours into ATGN+/-IPTG). Ph3/GFP/RFP images were acquired as shown in Fig.S9a. Among cells that displayed minimum two *ori1* foci (S-phase cells from Fig.S10), the number of *ori2*/cell was measured (from minimum 100 cells/condition) and means from 3 independent experiments were plotted for each condition. Error bars correspond to standard deviations. Student’s t-test: ns = P-value>0.01, ** = P-value<0.01.

**Figure 6:**
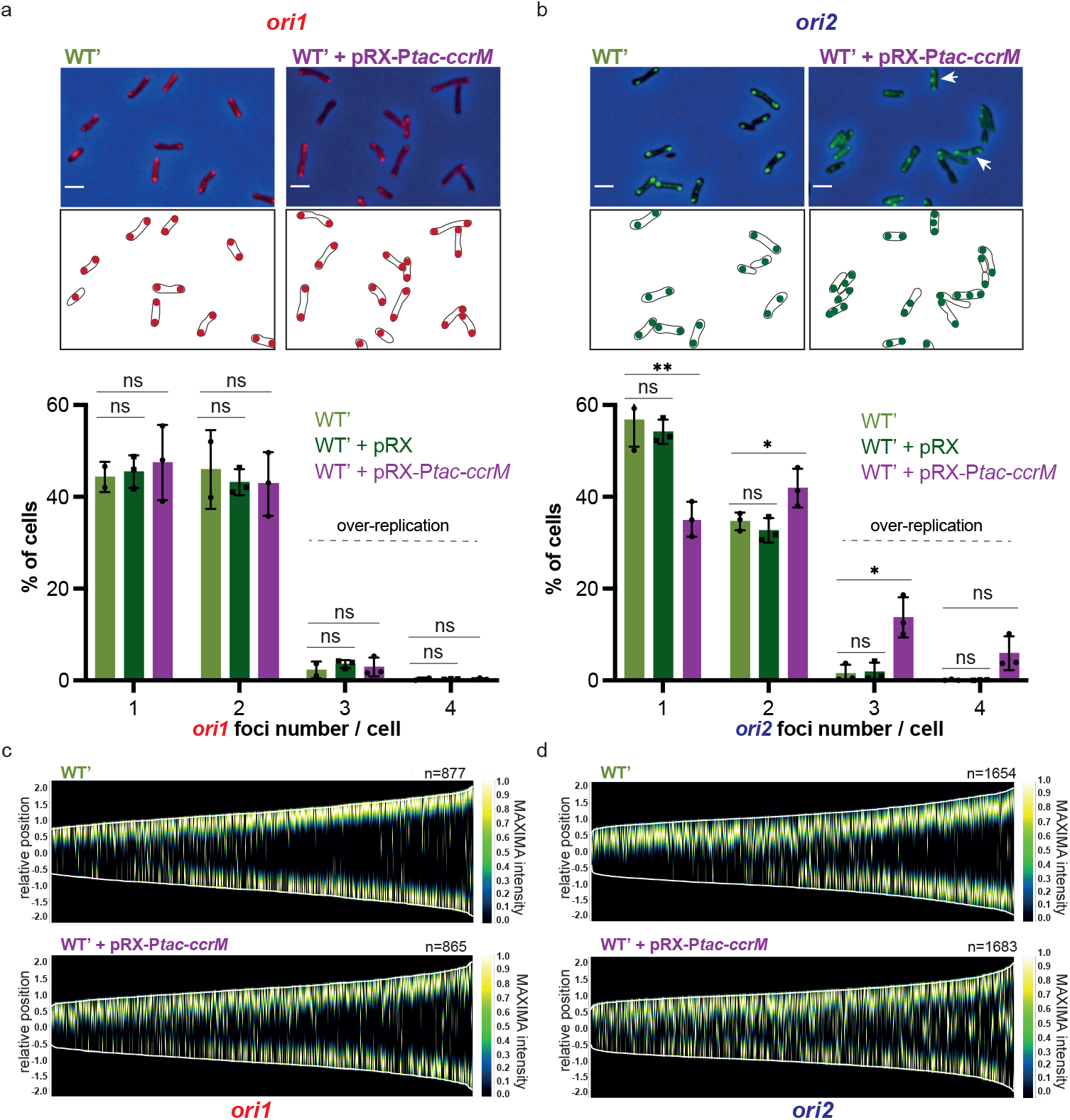
Overexpression of CcrM results in hyper-initiation events at *ori2* in cells cultivated in minimal medium. **(a)** and **(b):** Selected microscopy images of JC2656 (WT’ with *ori1/ygfp* reporter) cells (panel a) or JC2657 (WT’ with *ori2/ygfp* reporter) carrying or not the pRX-P*tac-ccrM* vector. Cells were cultivated over-night in ATGN+IPTG. Cultures were then diluted into ATGN+IPTG and grown exponentially for ∼6.5 hours (>15 hours into ATGN+IPTG). Upper panels: overlays of Ph3 and GFP images. Lower panels: schematics showing *ori1* (red color in **(a)**) or *ori2* (green color in **(b)**) subcellular localization in the cells imaged above. Under these microscopy images, plots show the proportion of cells with the indicated number of *ori1* (in **(a)**) or *ori2* (in **(b)**) foci for each strain (pRX is the empty control vector). Minimum 2 independent experiments (with a total of minimum 1000 cells) were done for each strain with each dot corresponding to an independent experiment and mean values were plotted with error bars corresponding to standard deviations. Student’s t-test: ns = P-value>0.05, * = P-value<0.05, ** = P-value<0.01**. (c)** and **(d):** Demographs showing the subcellular localization of *ori1* (panel c) or *ori2* (panel d) foci as a function of cell size (from images as shown in panels **(a)** and **(b)**). Relative position = 0 corresponds to mid-cell; only cells measuring from 1 to 4 μm-long were included into these demographs; n = number of cells used to construct each demograph.

How DNA methylation by CcrM promotes initiation from *ori2* in *A. tumefaciens* is not yet fully understood. Still, our data shows that the *repABC^Ch2^* operon, encoding all the elements required for replication initiation at *ori2* and *ori2* partitioning, is down-regulated when the *A. tumefaciens* genome becomes hypo-methylated (Fig.4a&b). Considering that the promoter region upstream of the *repABC^Ch2^* operon contains three GANTC motifs (Fig.1a right side and Tables S4&5), we propose that methylation of these motifs by CcrM promotes *repABC* transcription through an epigenetic mechanism that will be interesting to characterize in future studies. Noteworthy, the impact of CcrM on *repC^Ch2^* expression appeared as more striking than on *repA^Ch2^* and *repB^Ch2^* expression (Fig.4a&b). This observation suggests that a RepE^Ch2^ regulatory RNA may also play a role in controlling *repC^Ch2^* expression, as was previously shown for *repC^pTi^* control. Consistent with this proposal, we observed that the region between *repB ^Ch2^* and *repC ^Ch2^*, where a *repE ^Ch2^* gene is expected to be located, also carries three GANTC motifs that are efficiently methylated by CcrM (Fig.1a right side). Then, it is possible that DNA methylation by CcrM has a dual impact on *repC^Ch2^* expression by promoting its transcription through the methylation of the promoter upstream of *repABC^Ch2^* and through an inhibition of a putative *repE ^Ch2^* promoter (Fig.1a right side). A deeper analysis of our RNA-seq data indeed supported the existence of such anti-sense RepE RNAs located in the *repBC ^Ch2^* region and one of them is potentially repressed by CcrM (*Atu8080-3* in Tables S4&S6), although more experiments will be necessary to test this hypothesis more directly. Considering that the putative *ori2* inside *repC^Ch2^* also carries three conserved GANTC motifs (Fig.1a right side), it is also possible that *ori2* methylation by CcrM has a more direct impact on initiation at *ori2*.

Beyond the observed impact of CcrM on *repABC^Ch2^* expression, our transcriptome analysis revealed that DNA methylation by CcrM modulates the expression of many other *A. tumefaciens* genes. Gene Set Enrichment Analysis (GSEA) tests showed enrichments for COG categories J (translation/biogenesis), L (replication/recombination/repair) and N (cell motility) when comparing the transcriptome of CcrM-depleted cells with that of WT or CcrM-repleted cells (Fig.S11). The COG category N correlates with the motility/adhesion defects observed for CcrM-depleted cells (Fig.3), while the COG category L may correlate with up-coming replication issues, even if initiation from *ori2* still appeared as normal in most CcrM-depleted cells after 7h of growth in ATGN-IPTG (data not shown). Among genes that were significantly mis-regulated upon CcrM depletion, up to 75 may be directly regulated by CcrM-dependent methylation (Fig.4a and Table S5) and this “direct regulon” is strikingly different from previously described CcrM regulons in distantly-related bacteria ^11,21^ (Table S5). Moreover, the essential *ftsZ^At^* gene of *A. tumefaciens* (that has one GANTC motif in the 200 bp upstream of the ORF) was not significantly down-regulated in CcrM-depleted cells cultivated in complex media (Fig.4b), contrarily to what was previously observed in *C. crescentus* Δ*ccrM* mutant cells where FtsZ^Cc^ levels can become too limiting to sustain cell division and viability ^19,20^. Thus, even if *ccrM* is often (conditionally) essential in *Alphaproteobacteria ^19,22,23,35^*, the reason why *ccrM* can become essential appears to vary from one species to another. In *A. tumefaciens*, we propose that the essentiality of *ccrM* may instead be connected with its impact on *repC^Ch2^* expression and/or *ori2* methylation/activity to ensure genome maintenance.

Determining how promoter methylation by CcrM can impact gene expression in *A. tumefaciens* is an interesting perspective of this work. In *C. crescentus* and *B. subvibrioides*, GcrA is the most important epigenetic regulator that works together with CcrM to regulate gene expression ^18,21,36^. Tn-Seq experiments ^24^, and our own attempts to delete the *gcrA* gene of *A. tumefaciens* C58 strains (data not shown), indicate that *gcrA* is probably essential for cell viability/fitness. Here, we found that *gcrA* expression was significantly activated in cells with a hypo-methylated genome (Fig.4a&b, Table S5), suggesting that methylation of the *gcrA* promoter region may repress *gcrA* transcription (Fig.1a, right side). Consistent with this model, we found that this region carries five GANTC motifs (200 bp upstream of the ORF) (Fig.1a right side, Fig.S12a and Table S5), which is a strong over-representation compared to the average distribution of GANTC motifs on the *A. tumefaciens* chromosome (Fig.S6). In addition, we engineered a transcriptional reporter and thereby found that the *gcrA* promoter (250 bp upstream of ORF, containing 6 GANTC motifs) was significantly less active in WT or CcrM-repleted *A. tumefaciens* cells than in CcrM-depleted cells (Fig.S12b) and in *Escherichia coli* cells expressing an active *C. crescentus* CcrM MTase than in control cells expressing a catalytically inactive variant (FigS12c). If we now take into account the fact that the *A. tumefaciens gcrA* gene is located at a distance of ∼0.43 Mbp from *ori1* (Fig.1b), it is expected that it will switch from a fully-methylated to a hemi-methylated state only after a significant period of time following replication initiation at *ori1* (S-phase entry). This switch may then boost *gcrA* expression during the S-phase of the cell cycle (Fig.1a right side). If GcrA plays a role in the methylation-dependent regulation of *repC^Ch2^* and/or *repE^Ch2^* (Fig.1a, right side), it could possibly act as a sophisticated timer coordinating initiation at *ori2* with replication progression from *ori1* to ensure robust genome maintenance over generations. Future studies should then focus on testing whether GcrA is, indeed, the main epigenetic regulator working together with CcrM in *A. tumefaciens* to shed light on (epigenetic) regulatory networks that primitive organisms with complex genomes developed to coordinate genome replication from multiple origins the way more complex organisms also do.

## METHODS

### Bacterial growth conditions

*E. coli* strains were grown using standard growth conditions in/on Luria-Bertani (LB: 1% tryptone, 0.5% yeast extract, 1% NaCl, pH 7) +/- bacto-agar (1.5%) medium at 37^ο^C. Antibiotics were added at the following concentrations when needed (liquid/solid media): kanamycin 30/50 μg/mL, oxy-tetracycline 12/12 μg/mL, ampicillin 50/100 μg/mL. *A. tumefaciens* strains were grown at 28°C in LB, Yeast Extract Beef ^37^ (YEB from fisher scientific Bioworld 306270311: 0.5% tryptone, 0.1% yeast extract, 0.5% nutrient broth, 0.5% sucrose, 0.049% MgSO_4_-7H_2_O, pH 7.2) or AT minimal medium (without exogenous iron) ^38^ with 0.5% glucose (ATGN) or 5% sucrose (ATSN). Agar plates and soft-agar plates were prepared with 1.35% (ATGN), 1.2% (ATSN) or 0.25% (soft) bacto-agar, respectively. Antibiotics were added at the following concentrations when needed (liquid/solid media): kanamycin 150/300 μg/mL in all media, gentamycin 100/200 in ATGN/ATSN, oxy-tetracycline 3/3 μg/mL in ATGN. When indicated, IPTG was added at a final concentration of 1mM.

### Bacterial strains, plasmids and oligonucleotides

Bacterial strains, plasmids and primers used in this study are listed in Supplementary Tables S1, S2 and S3, respectively. The genomes of the *A. tumefaciens* C58 derivatives (from ^29^) used in this study were analyzed by PCR as described before ^28^ and this analysis showed that these strains are so-called “C58 fusion strains” with a unique dicentric chromosome (Fig.S13). This feature was subsequently verified when re-assembling the genomes of the WT/JC2307 strains from whole-genome SMRT-seq data.

### Construction of plasmids and strains

Plasmids were constructed using standard DNA cloning techniques and the inserts of all constructs were verified by Sanger sequencing (Microsynth AG or Sigma AG). *E. coli* transformations were carried out using standard methods. When necessary, replicating or integrative plasmids were introduced into *A. tumefaciens* cells by conjugation or electroporation ^29^. *E. coli* S17.1 *λpir* or TOP10 strains were used for conjugation and cloning procedures, respectively.

Construction of pUC-miniTn7TGMPtac-ccrM: The *ccrM* (*Atu0794*) ORF of the *A. tumefaciens* C58 strain was amplified using primers MS9 and MS10, digested with NdeI/BglII and ligated into NdeI/BamHI-digested pUC-miniTn7TGMPtac-HA.

Construction of pNPTS138-ΔccrM: The 500 bp sequences upstream and downstream of the *A. tumefaciens* C58 *ccrM* ORF were amplified using primer pairs MS1/MS2 and MS3/MS4, and digested by HindIII/BamHI and BamHI/NheI, respectively. Both fragments were then ligated into HindIII/NheI-digested pNPTS138.

Construction of pRX-P*tac-ccrM:* The P*tac-ccrM* construct was amplified from pUC-miniTn7TGMPtac-ccrM using primers MS40 and MS41, digested by EcoRI/BglII and cloned into EcoRI/BglII-digested pRXMCS-2.

Construction of the JC2291 strain: The pUC-miniTn7TGMPtac-ccrM plasmid was introduced into the JC2141 strain by electroporation with the pTNS3 helper plasmid. Insertion of P*tac-ccrM* into the JC2291 genome was verified using primers Tet-Forward and Tn7R109 as described before ^29^.

Construction of the JC2307 strain: The pNPTS138-ΔccrM plasmid was introduced into the JC2291 strain by conjugation (kanamycin-resistant *A. tumefaciens* colonies were selected on ATGN+IPTG as *E. coli* S17.1 λpir does not grow on ATGN ^39^). Integration at the native *ccrM* locus of the JC2291 chromosome by homologous recombination was verified by checking sucrose-sensitivity and by colony-PCR using primers MS4 and MS17. The resulting strain was then cultivated over-night in ATGN+IPTG medium (without antibiotics) before plating onto ATSN+IPTG medium (plasmid re-excision step). Kanamycin-sensitive and sucrose-resistant colonies were then selected onto ATSN+IPTG+/-kanamycin media. We then screened for Δ*ccrM* colonies by colony PCR using primers MS1 and MS4.

Construction of JC2656/JC2657/JC2777/JC2660/JC2661: pWX963 or pWX967 were introduced into JC2141 or JC2307 by conjugation using a kanamycin-resistance selection on ATGN+IPTG.

Construction of JC2277/JC2836: JC2657 and JC2661 were cultivated over-night in ATGN+/- IPTG medium (without antibiotics, for pWX963 re-excision step) before plating onto ATSN+/-IPTG medium. Kanamycin-sensitive and sucrose-resistant colonies were then selected onto ATGN+/-IPTG. We then screened for colonies that kept the *ori2/ygfp* reporter by fluorescence microscopy. The pWX995 plasmid (donor of *ori1/mcherry* reporter) was then introduced into these two kanamycin-sensitive by conjugation using a kanamycin-resistance selection on ATGN+IPTG.

### Microscopy

To visualize and analyze cell morphology, cells were fixed using a 5X-fix solution (150mM NaPO_4_, 12.5% formaldehyde at pH7.5) and stored at 4^°^C before being imaged on 0.5X PBS (phosphate-buffered saline) and 1% agarose pads using a Plan-Apochromat 100X/1.45 oil Ph3 objective on an AxioImager M1 microscope (Zeiss) with a cascade 1K EMCCD camera (Photometrics) controlled by the VisiView 7.5 software. For phase-contrast and fluorescence microscopy, live cells were immobilized onto 0.5X PBS + 1% agarose pads and imaged using the same microscope system as described above. To analyze the subcellular localization and the number of fluorescent origin foci, and to construct demographs, image analyses were performed using the Fiji 2.3.0 software with the MicrobeJ plugin (default parameters) ^40^.

### gDNA preparation

Genomic DNA samples were prepared from 1-2mL of bacterial cultures. Cells were pelleted and immediately frozen in liquid nitrogen prior to conservation at -80^°^C. gDNA was extracted using an isopropanol-ethanol purification kit (Puregene Yeast/Bact. Kit B from Qiagen) and following the manufacturer’s protocol including a 60-minute RNase A treatment. gDNA was rehydrated in H_2_O. gDNA sample quality and quantity were assessed using a Nanodrop spectrophotometer before storage at -20^°^C.

### SMRT-sequencing and analyses

High molecular weight gDNA was sheared with the Megaruptor 3 (Diagenode) to obtain 10- 15 kbp fragments. After shearing, the DNA size distribution was checked using a Fragment Analyzer (Agilent Technologies). Multiplexed SMRTbell libraries were prepared from 365ng of sheared DNA with the PacBio SMRTbell Express Template Prep Kit 2.0 (Pacific Biosciences) according to the manufacturer’s recommendations (protocol 101-696-100, v07). A final size selection step was performed with Pacific Biosciences Ampure beads to remove libraries smaller than 3 kbp in size. The resulting libraries were pooled and sequenced over a 15H movie length on a single PacBio Sequel II SMRT cell 8M using the Binding kit version 1.0 and the Sequencing kit version 2.0 (Pacific Biosciences). Sequences assembly was performed with SMRTlink software suite version 9.0.0.92188 (Pacific Biosciences, USA). To analyze the methylome of our strains, the raw data were processed using the SMRTlink version 10.2.0.133434 (Pacific Biosciences, USA). The tool named "Base Modification Analysis Application" was then used to detect modified DNA bases and to identify methylated DNA motifs. The sequencing reads were mapped on the assembled genome sequences of the dicentric C58 strain and analyzed using default parameters with the "Find Modified Base Motifs" option. For each analyzed sample, the tool produced a table listing the genomic positions at which a base modification is detected, as evidenced by the deviation of the inter-pulse duration from the expected base incorporation time. The tool also generated a list of motifs that are present in more than 30% of the occurrences of that motif around detected modified bases. A perl script was then written to extract the coverage, IPD value and score values for each GANTC motif occurrence in the whole genome; an IPD value of 1.0 was attributed to GANTC motifs where the software did not detect a methylation event.

### Soft-agar motility assays

Swim plate assays were conducted as described previously ^38^ with minor adaptations (see Fig.3a legend). Swimming area were measured from images using the ImageJ 2.3.0 software.

### Biofilm/surface attachment assays

Biofilm assays were conducted as described previously ^38^ with minor adaptations. Briefly, 4 times 3mL of each culture diluted to an OD_600_∼0.05 were introduced into 4 wells containing a size-adjusted, vertical polyvinyl chloride (PVC) coverslip. 12-well polystyrene plates were incubated at room-temperature for 72 hours into a sealed container containing a smaller recipient with 200mL of saturated sulfate solution (120g/L). After incubation, the coverslips were removed from wells and immersed into deionized H_2_O to remove planktonic cells. Dried coverslips were then immersed into 1mL of a 0.1% crystal violet (CV) solution for 5 minutes prior to rinsing with deionized H_2_O and careful drying. For each assay, one coverslip was kept for imaging, while the other three were each immersed into 1mL of a 33% acetic acid solution. The A_600_ of the solubilized CV and the OD_600_ of the cultures from each well were then measured. The biofilm score was calculated as the A_600_ divided by the OD_600_.

### RNA extraction

RNA samples were prepared from 4-5 mL of bacterial cultures. Cells were pelleted and immediately frozen in liquid nitrogen prior to conservation at -80^°^C. RNA were extracted using an RNeasy Mini-Kit from Qiagen and following the manufacturer’s protocol and including a DNase I (RNase-free DNases set from Qiagen) treatment. Samples were additionally treated with a TURBO DNA-free kit from Invitrogen following the manufacturer’s protocol. Absence of DNA contaminations were verified by standard PCR. Quality and quantity of RNA samples were verified on an agarose gel and using a Nanodrop spectrophotometer before storage at -80^°^C.

### RNA-sequencing and analyses

RNA quality was assessed using a Fragment Analyzer (Agilent Technologies) and all RNAs had RQNs between 9.7 and 10. RNA-seq libraries were prepared from 800 ng of total RNA with the TruSeq Stranded mRNA Prep reagents (Illumina) using a unique dual indexing strategy, and following the official protocol automated on the Sciclone liquid handling robot (PerkinElmer). The polyA selection step was replaced by an rRNA depletion step with the QIAseq FastSelect - 5S/16S/23S bacterial rRNA removal kit (Qiagen). Libraries were quantified by a fluorometric method (QubIT, Life Technologies) and their quality assessed using the Fragment Analyzer. Sequencing was performed on the Illumina HiSeq 4000 v4 SR flow cell with the v4 HiSeq 3000/4000 SBS Kit reagents for 150 cycles. Sequencing data were demultiplexed using the bcl2fastq2 Conversion Software (version 2.20, Illumina). Sequences matching to ribosomal RNA sequences were removed with fastq_screen (v. 0.11.1)^41^. Remaining reads were further filtered for low complexity with reaper (v. 15-065) ^42^. Reads were aligned against the *Agrobacterium fabrum C58 ASM9202v1* genome using STAR ^43^ (v. 2.5.3a). The number of read counts per gene locus was summarized with htseq-count ^44^ (v. 0.9.1) using a custom *Agrobacterium fabrum C58 ASM9202v1* gene annotation including 7 small RNA genes listed in Table S6. Quality of the RNA-seq data alignment was assessed using RSeQC ^45^ (v. 2.3.7). Counts per gene table was used for statistical analysis in R (R version 4.1.0). Genes with low counts were filtered out according to the rule of 1 count per million (cpm) in at least 1 sample. Library sizes were scaled using TMM normalization (EdgeR package ^46^ version 3.34.0) and log-transformed with limma cpm function (Limma package ^47^ version 3.48.0). Differential expression was computed with limma by fitting the samples into a linear model and performing comparisons with moderated t-test. Global p-value adjustment with Bonferroni-Hotchberg method was used for all comparisons. Genes with an adjusted P-value<0.01 and a minimum fold-change of 2 were considered as significantly mis-regulated. Volcano plots showing differentially expressed genes were made in R (v4.2.1) by plotting the log2(fold change) against the -log2(adjusted P-value) for this comparison.

### qRT-PCR experiments and analyses

Primers were chosen based on secondary structure predictions using the Mfold web server for nucleic acid folding and hybridization prediction (default parameters, except “folding temperature” set to 60^°^C) ^48^. Primer pairs were then checked by standard PCR on *A. tumefaciens* C58 gDNA (1ng/μL) to verify that they gave a single amplification product of the expected size. Primer pairs were then checked again by qPCR (4 times 4-fold serial dilutions) to check efficiency). 700 ng of RNA samples were retrotranscribed using the Verso cDNA synthesis kit (Thermoscientific) for 30 minutes at 42^°^C into a final volume of 20μL and following the manufacturer’s protocol. The Verso reverse transcriptase was then inactivated by a 2-minute incubation at 95^°^C before cDNA samples were stored at -20^°^C. For each qPCR assays, 2.4μL of cDNA samples diluted 1:20 in Tris 10mM (pH 8.5) were used as templates. The qPCR reaction mix contained 0.3μM of each primer into 10μL of Power SYBR Green PCR Master Mix (Applied Biosystems). The 284-well plate was filled using an Evo® TECAN robot and transferred into a qPCR machine (Thermofisher - QuantStudio6) with automated threshold calculations (Software QuantStudio Q6_v1.6). Cycling: 10 s at 95°C, 15 s at 95°C and 1 minute at 60°C for 40 cycles, 15 seconds at 95°C. Differences in gene expression were evaluated based on stable internal gene controls chosen from the RNAseq data: *purH (Atu_2823,* bifunctional purine biosynthesis protein) for ATGN +/- IPTG 7h conditions or *yidC (Atu_0384,* membrane protein insertase YidC) for YEB +/- IPTG 5.5h conditions. The Delta-Delta Ct method ^49^ was used to estimate relative mRNA levels from averages calculated from three technical replicates. Three biological replicates were used for each strain/condition tested.

### Statistics and reproducibility

Statistical methods and sample sizes (n) are indicated in figure legends for each experiment. Statistical analyses were done using Excel, GraphPad-PRISM or the R softwares.

## DATA AVAILABILITY

Data that support the findings of this study are available from the corresponding author upon request.

## Supporting information

GlimmaVolcanoPlot

## ACKNOWLEDGEMENTS

We thank Pamela Brown, Xindan Wang, Patrick Viollier, Gaël Panis and Albert Jeltsch for gifts of strains/plasmids/antibodies. We also thank Hannes Richter, Jordan Vacheron, Clara Heiman, Garance Sarton, Jessica Burnier and Gaël Close for technical help during statistical/bioinformatic/microscopy analyses. We thank Jacqueline Masternak and Laurent Casini for several plasmid constructions, together with Noémie Matthey and Giorgia Wennubst for discussions during the project and feedback on the manuscript. This work was funded by the Swiss National Science Foundation grants 31003A_173075 and 310030_204822 to J.C.

