## Supplementary Information for "DNA methylation by CcrM contributes to genome maintenance in the *Agrobacterium tumefaciens* plant pathogen"

**CONTENT:**

Page 2: SUPPLEMENTARY FIGURES WITH LEGENDS

Page 13: SUPPLEMENTARY TABLES WITH LEGENDS

Page 18: SUPPLEMENTARY METHODS

Page 19: SUPPLEMENTARY REFERENCES

**SUPPLEMENTARY FIGURES WITH LEGENDS:**

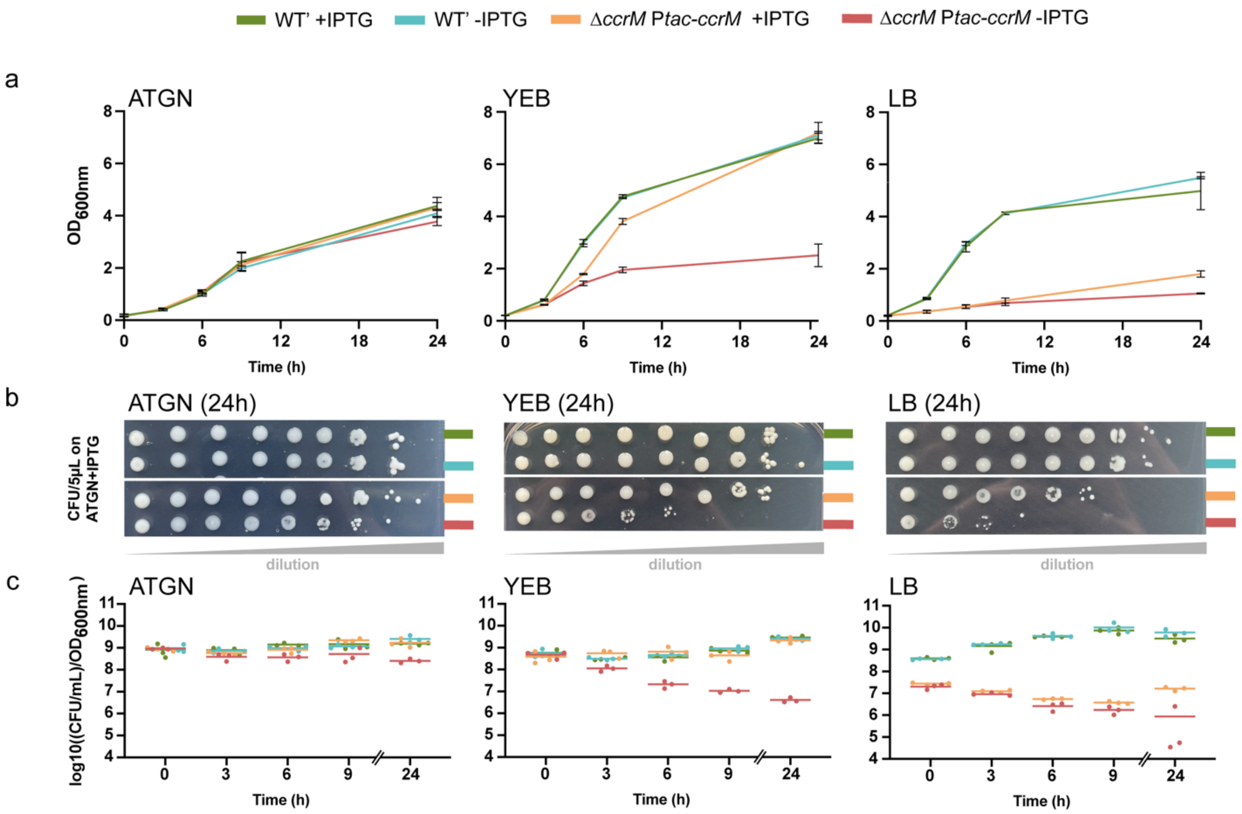


**Figure S1: CcrM-depleted cells loose viability over time when cultivated in complex media. (a)** JC2141 (WT') and JC2307 (*ΔccrM* P*tac-ccrM*) cells were pre-cultivated into ATGN+IPTG medium and cultures were diluted back into the indicated media +/-IPTG in test tubes. Growth was evaluated over-time measuring the $OD_{600}$. **(b)** After 24h of growth from (a), 5µL samples of each culture were collected and spotted onto solid ATGN+IPTG medium (= first undiluted sample on the left of each plate lane). Serial 10-fold dilutions of each culture were also plated (from left to right sides of images) to compare the capacities of cells to form colony forming units (CFU) on permissive plates (JC2307 grows relatively well onto ATGN+IPTG medium as can be seen in Fig.2) depending on the liquid medium into which cultures were grown. **(c)** Quantitative comparison of CFU using three biological replicates for each condition/strain (one example of each is shown in panel (b)).

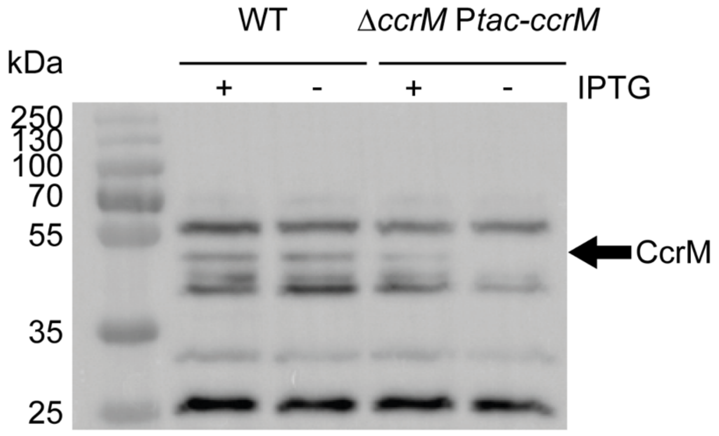


**Figure S2: Immunoblot experiments showing that CcrM is depleted in JC2307 cells cultivated in ATGN-IPTG for 7 hours.** JC2140 (WT) and JC2307 (*ΔccrM* P*tac-ccrM*) cells were cultivated into ATGN+/-IPTG as described in Fig.S1a and samples were collected at the 7-hour time point. Proteins were separated on a 12% SDS-PAGE gel and transferred onto a PVDF membrane (Millipore). Immunoblotting was performed following a standard protocol [1] and using a rabbit anti-CcrM$^{At}$ serum (generous gift from the Viollier lab, University of Geneva) diluted 1:5000 and a goat anti-rabbit (A9169 from Sigma) diluted 1:30000. The black arrow indicates which protein most likely corresponds to CcrM (depleted when IPTG is removed and fitting the expected molecular weight of CcrM$^{At}$ of ~42.2 kDa). Other detected proteins can be used as loading controls.

***→ Also note that the exact same ATGN+/-IPTG culture samples as described here were used for the SMRT-Seq (Fig.1b and Fig.S6 below), RNA-Seq (Fig.4 and Fig.S7 below) and HinfI-based (Fig.S5 below) experiments described in this study.***

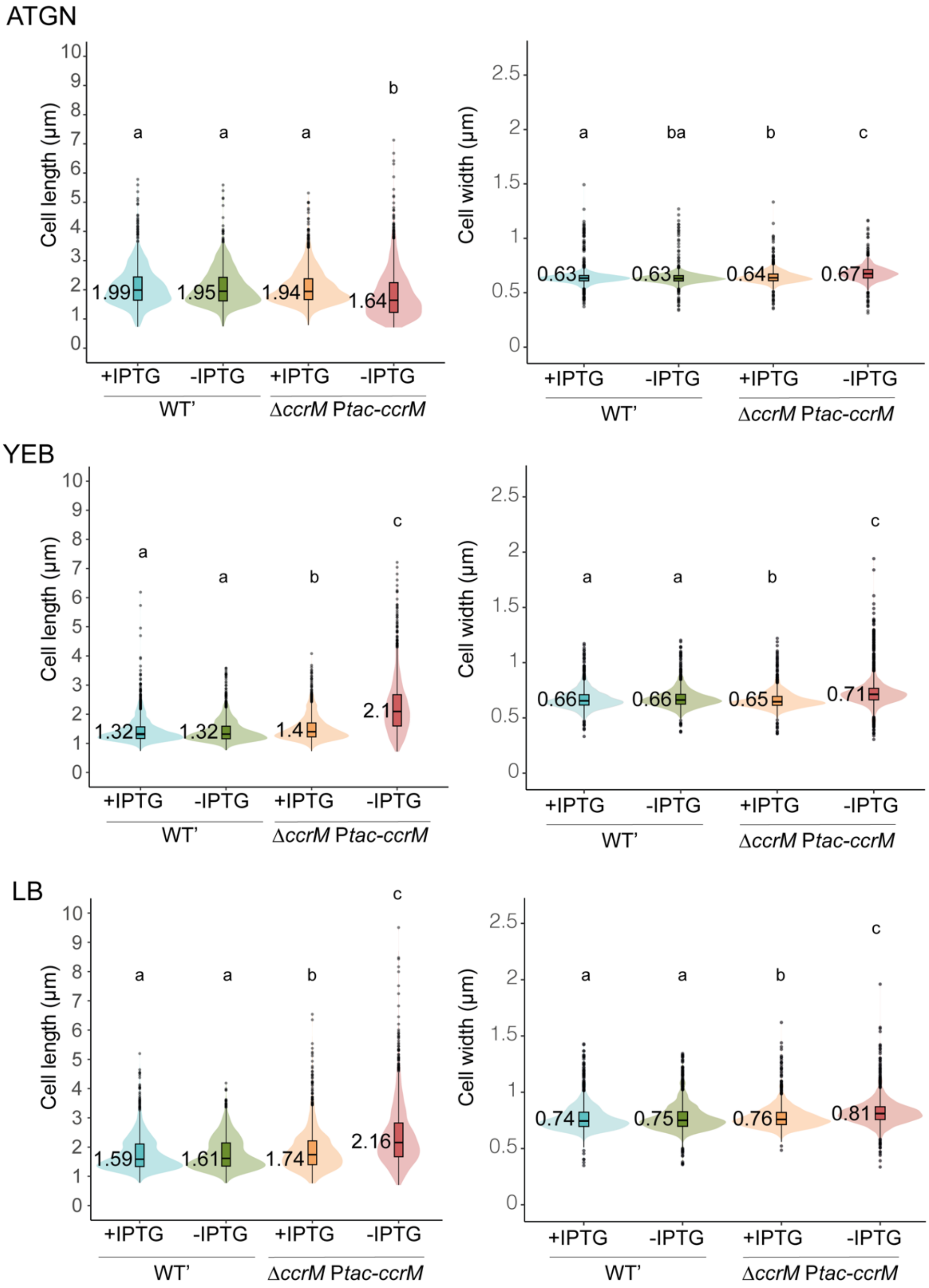


**Figure S3: Microscopy image analyses comparing the length/width of JC2141 (WT') and JC2307 (*ΔccrM* P*tac-ccrM*) cells cultivated in different media.** Strains were cultured and imaged as described in Fig.2c (24 hours of growth in the indicated media). The length and the width of 1500 cells for each strain/condition were then measured from phase-contrast images using the Fiji 2.3.0 software with the MicrobeJ plugin [2]. Written values indicate the median cell length or width in the cell population. Boxplots include the 25-75 percentiles. The potential significance of differences was evaluated for each plot using a Kruscal-Wallis test with Bonferroni corrections. Note that differences in cell widths are globally subtle even if 2-3 statistically different groups (a, b or c) can be distinguished (P-value<0.001) for each plot using this test.

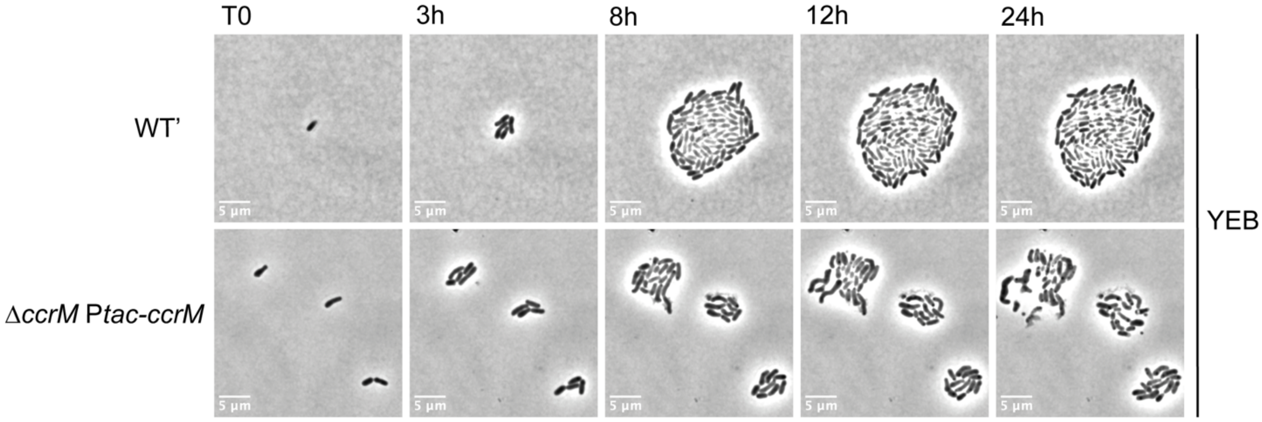


**Figure S4: Time-lapse microscopy images of JC2141 (WT') and JC2307 *(ΔccrM* P*tac-ccrM*) cells cultivated in YEB without IPTG.** Strains were pre-cultured to stationary phase in YEB+IPTG and then diluted back into YEB+IPTG. Once cells reached exponential phase again, they were washed and resuspended into YEB-IPTG to be spotted onto a YEB-IPTG agarose (1%) pads (T0) for phase-contrast microscopy. Cells were imaged every 15 minutes for 24 hours using a 100x/1.40 inverted oil-immersion objective on a Leica DMi8 microscope with a sCMOS DFC9000 (Leica) camera and SOLA light engine (Lumencor). Representative images of developing micro-colonies are shown in this figure. Movies showing microcolony growth over time are also available: Movie S1 (WT') and Movie S2 (*ΔccrM* P*tac-ccrM).*

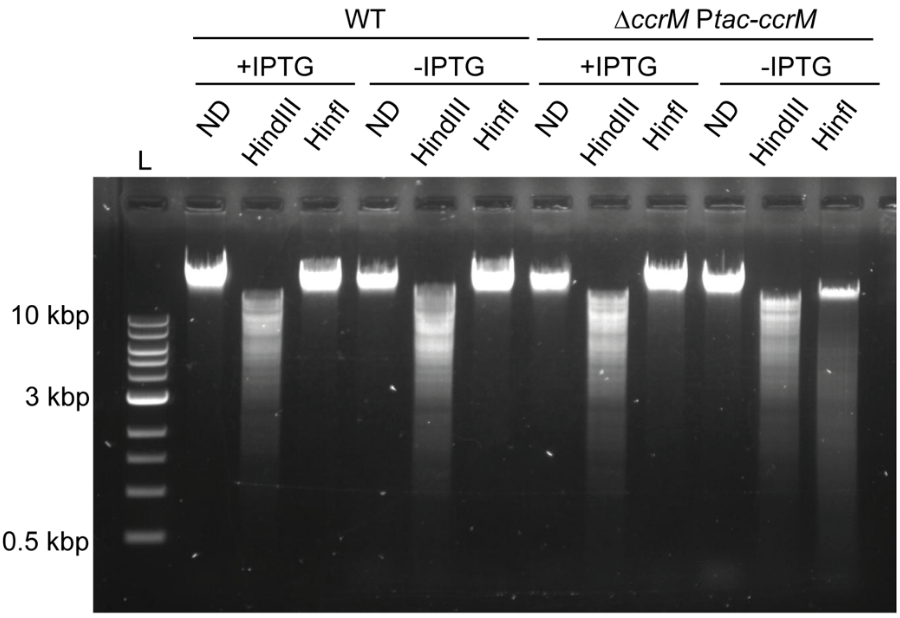


**Figure S5: HinfI-based assay indicating that the genome of CcrM-depleted cells becomes hypo-methylated.** JC2140 (WT) and JC2307 (*ΔccrM* P*tac-ccrM*) cells were cultivated into ATGN+/-IPTG as described in Fig.S1a and samples were collected at the 7-hour time point for gDNA extraction. 2 µg of gDNA was kept non-digested (ND) or digested with HindIII (digestion control, independent of DNA methylation state) or HinfI (cuts only at non-methylated GANTC motifs) for 4 hours at $37^0$C. gDNA fragments were then separated by gel electrophoresis using a TAE 1X with 0.8% agarose gel. L: DNA ladder, with the white arrow showing a DNA band at 3kbp.

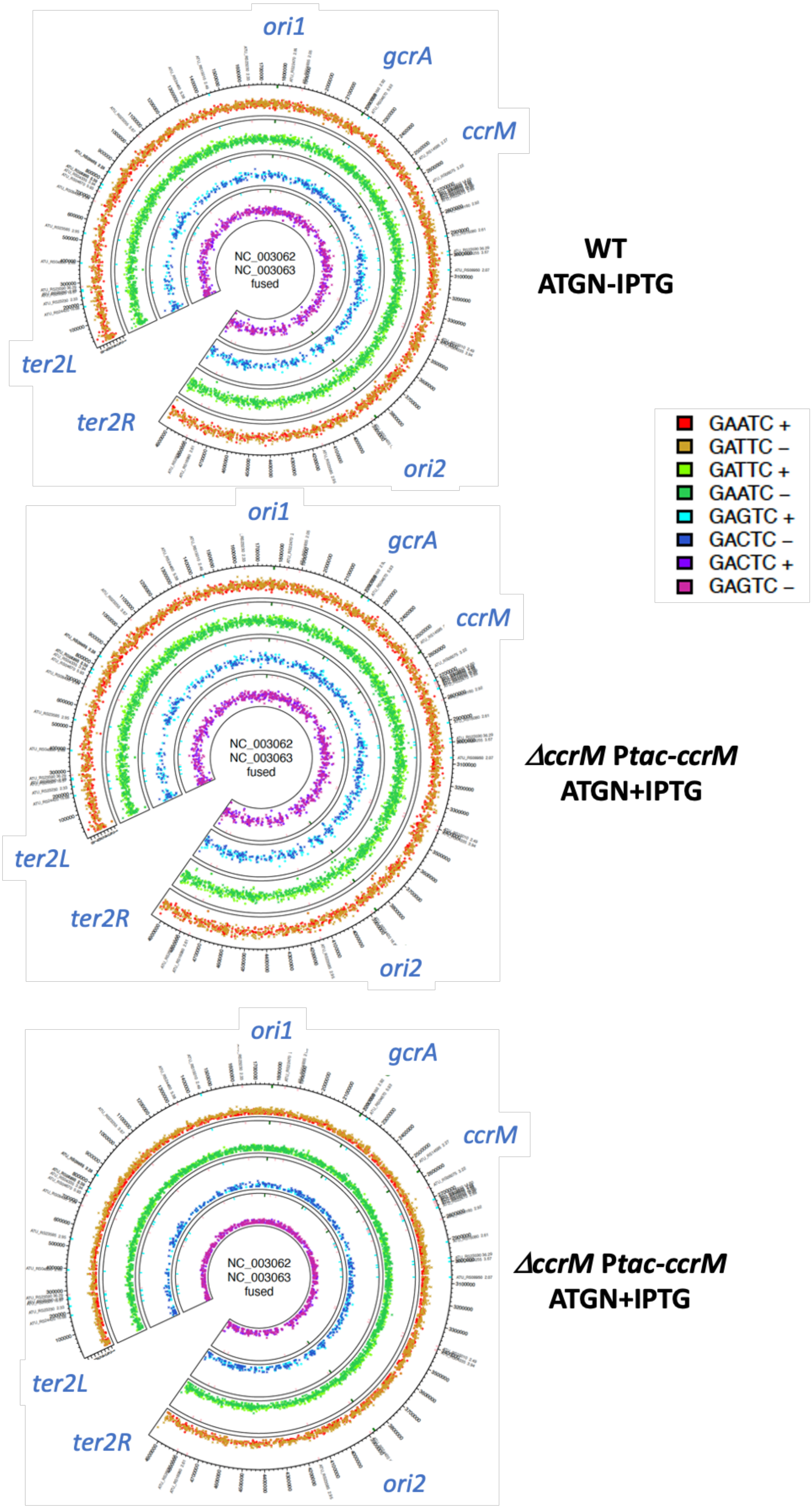
ori1
gcrA
ccrM
NC_003062
NC_003063
fused
ter2L
ter2R
ori2
WT
ATGN-IPTG
ori1
gcrA
ccrM
NC_003062
NC_003063
fused
ter2L
ter2R
ori2
ΔccrM Ptac-ccrM
ATGN+IPTG
ori1
gcrA
ccrM
NC_003062
NC_003063
fused
ter2L
ter2R
ori2
ΔccrM Ptac-ccrM
ATGN+IPTG
GAATC +
GATTC –
GATTC +
GAATC –
GAGTC +
GACTC –
GACTC +
GAGTC –

**Figure S6 (previous page)**: **Distribution and IPD ratio (y axis) of individual GANTC motifs (+ or – strands) on the dicentric linear chromosome (from *ter2L* to *ter2R*) of JC2140 and JC2307 cells cultivated in ATGN+/-IPTG for 7 hours.** Cells were cultivated as described in Fig.S1a and samples were collected at the 7-hour time point for gDNA extraction followed by SMRT-Seq as also described in Fig.1b. The same y scale (for IPD ratio) was used in all three schematics.

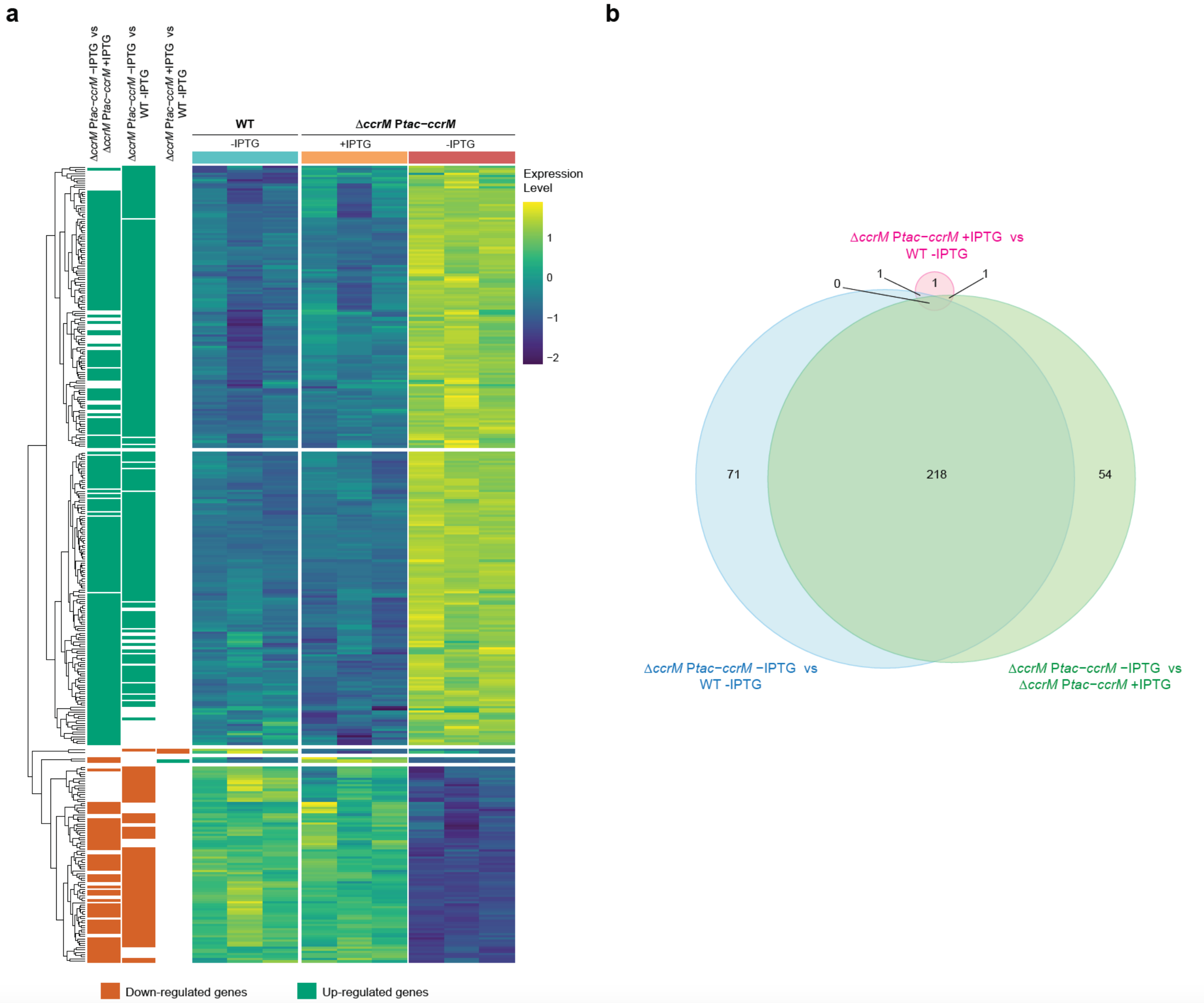


**Figure S7: Comparison of the transcriptome of JC2140 and JC2307 cells cultivated in ATGN +/- IPTG for 7 hours.** Cells were cultivated into ATGN+/-IPTG as described in Fig.S1a and samples were collected at the 7-hour time point for RNA-seq analyses. **(a)** Heatmap of significantly mis-regulated genes (adj.P-value<0.01 and min 2-fold change), ordered by genotypes: WT (JC2140) -IPTG, Δ*ccrM* P*tac-ccrM* (JC2307) +IPTG and Δ*ccrM* P*tac-ccrM* -IPTG. Expression values for each gene were scaled and centered. Colors on the left-hand side indicate if genes are down-regulated (orange) or up-regulated (green). **(b)** Venn diagram showing the number of significantly mis-regulated genes from panel **(a)**. Each circle corresponds to one comparison and the circle size partially reflects the number of mis-regulated genes in the given comparison.

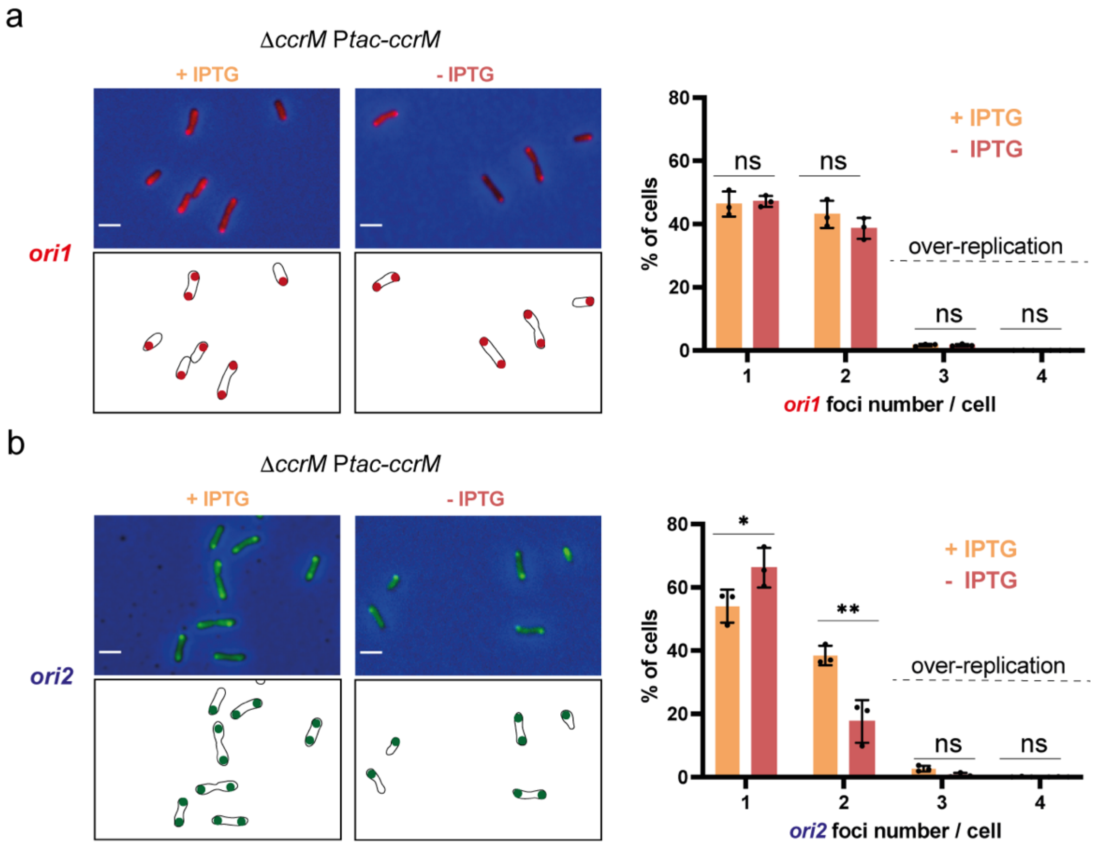


**Figure S8: Localization and number of *ori1* or *ori2* (single *ori* labelling) in CcrM-repleted and CcrM-depleted cells cultivated in ATGN medium. Left side:** Selected microscopy images of JC2660 (*ΔccrM* P*tac-ccrM* with *ori1/ygfp* reporter) cells in panel **(a)** or JC2661 cells (*ΔccrM* P*tac-ccrM* with *ori2/ygfp* reporter) cells in panel **(b)** cultivated over-night in ATGN+/-IPTG. Cultures were then diluted into ATGN+/-IPTG and grown exponentially for ~6.5 hours (>15 hours into ATGN+/-IPTG). Upper panels: overlays of Ph3 and GFP images. Lower panels: schematics showing *ori1* (red color in **(a)**) or *ori2* (green color in **(b)**) subcellular localization in the cells imaged above. **Right side:** Quantification of *ori1* or *ori2* number per cell from multiple microscopy images (including minimum 1000 cells). Mean values from 3 independent experiments were plotted for each strain/condition. Error bars correspond to standard deviations. Student's t-test: ns: P-value>0.05, *: P-value < 0.05, **: P-value<0.01.

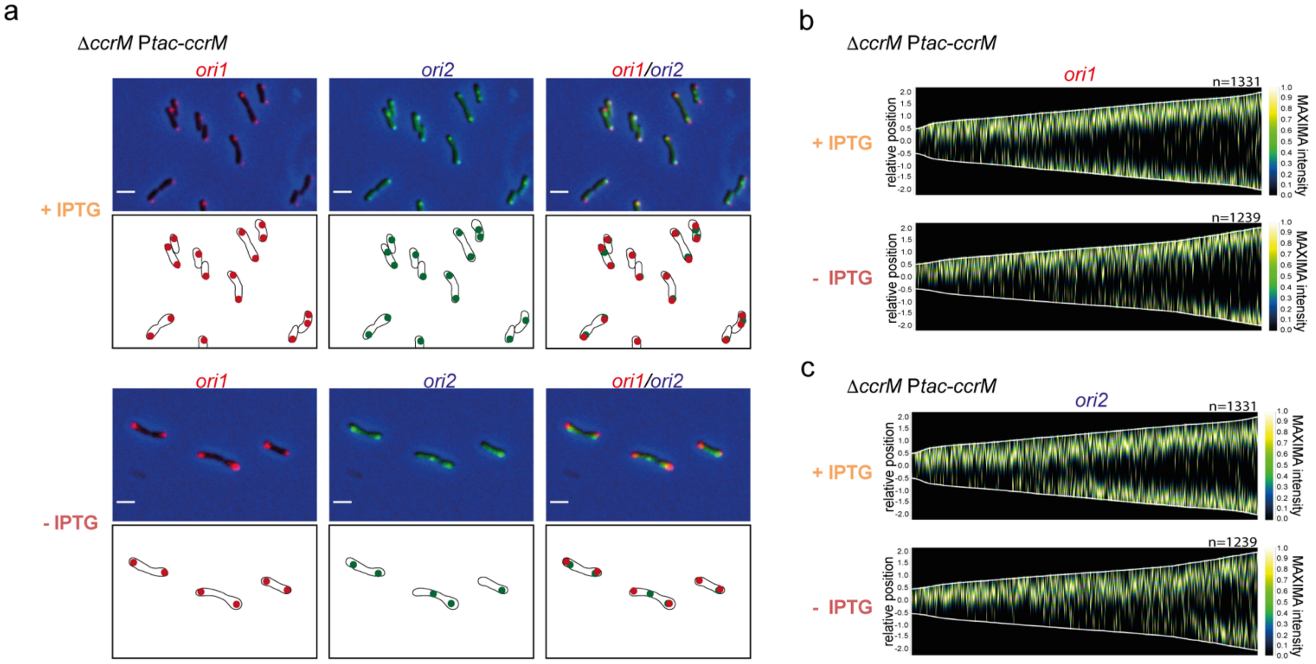


**Figure S9: Localization and number of *ori1* and *ori2* (double *ori* labelling) in CcrM-repleted and CcrM-depleted cells cultivated in ATGN medium. (a)** Selected microscopy images of JC2836 (*ΔccrM* P*tac-ccrM* with *ori1/mcherry* and *ori2/ygfp* reporters) cells cultivated over-night in ATGN+/-IPTG. Cultures were then diluted into ATGN+/-IPTG and grown exponentially for ~6.5 hours (>15 hours into ATGN+/-IPTG). Upper panels: overlays of Ph3 and GFP and/or RFP images. Lower panels: schematics showing *ori1* (red color) or *ori2* (green color) subcellular localization in the cells imaged above. **(b)** Demographs showing the subcellular localization of *ori1* and *ori2* foci as a function of cell size (same conditions as in panel **(a)**). Relative position = 0 corresponds to mid-cell. Only cells measuring from 1 to 4 μm-long were included into these demographs. n: number of cells used to construct each demograph.

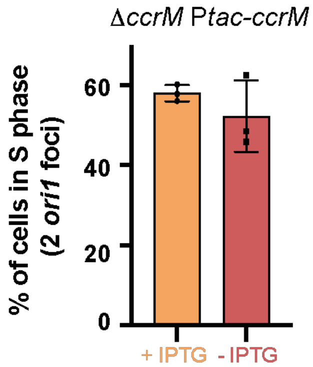


**Figure S10: Proportion of S-phase cells in JC2836 (*ΔccrM* P*tac-ccrM* with *ori1/mcherry* and *ori2/ygfp* reporters) cells.** Cells were cultivated over-night in ATGN+/-IPTG. Cultures were then diluted into ATGN+/-IPTG and grown exponentially for ~6.5 hours (>15 hours into ATGN+/-IPTG) as described in Fig.S9. The number of *ori1* (red foci)/cell was measured as a *proxi* to evaluate the % of S-phase cells in each population. Three independent cultures were used for each condition (minimum 500 cells/condition). Mean % of cells were plotted. Error bars correspond to standard deviations. A student's t-test indicates that the difference is not significant (P-value = 0.19).

| COG category | *ΔccrM* P*tac-ccrM* +IPTG versus WT-IPTG | *ΔccrM* P*tac-ccrM* -IPTG versus WT-IPTG | *ΔccrM* P*tac-ccrM* -IPTG versus *ΔccrM* P*tac-ccrM* +IPTG |
|---|---|---|---|
| **J (translation/biogenesis)** | NS | P-value=0.0115 | P-value=0.0115 |
| **L (replication/recombination/repair)** | NS | P-value=0.0115 | P-value=0.0115 |
| **N (cell motility)** | NS | P-value=0.0153 | NS |

**Figure S11: COG categories J, L and N are significantly over-represented among genes that are mis-regulated in CcrM-depleted cells compared to CcrM-repleted or WT cells**. Strains, growth conditions and RNA samples are the same as described in Fig.S7. A Gene Set Enrichment Analysis (GSEA) was performed using the R package BOG (Bacterium and virus analysis of Orthologous Groups) [3] to identify over-represented COG categories among genes that were significantly mis-regulated (FC>2 and adjusted P-value<0.01 in Table S4) when comparing the indicated strains/conditions. P-values were adjusted for multiple testing. NS = no significant enrichment.

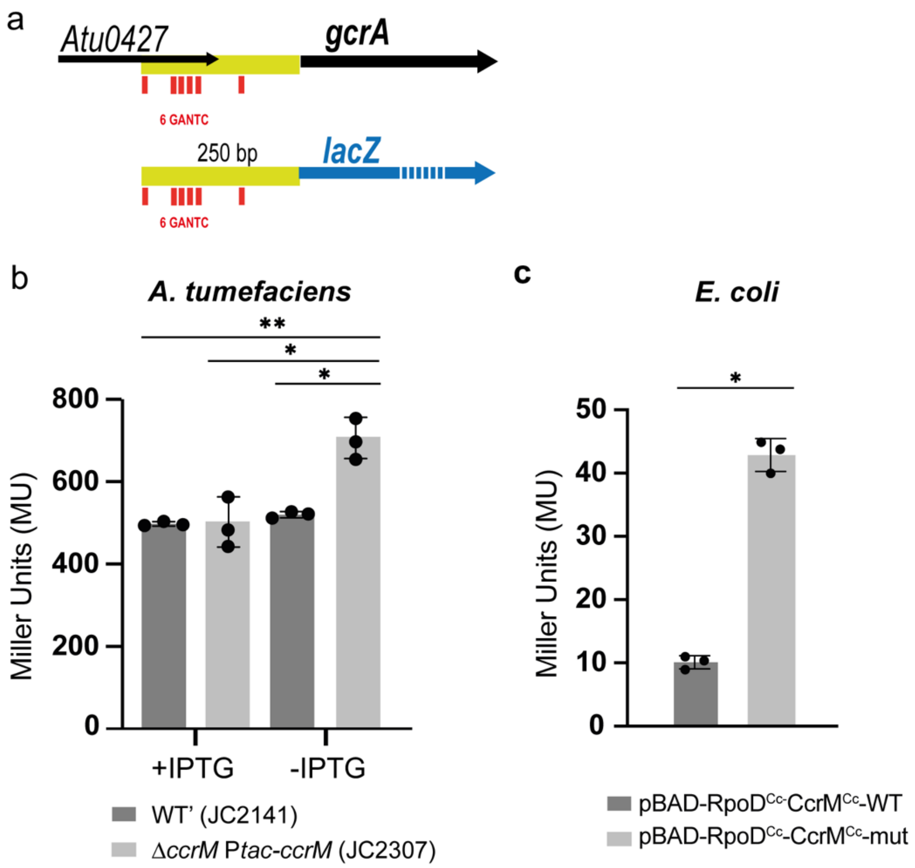


**Figure S12: CcrM represses the *A. tumefaciens gcrA* promoter in *A. tumefaciens* and *E. coli* cells.** **(a)** Map of the *gcrA (Atu0426)* promoter region (yellow) and position of GANTC motifs (red) in this region on the *A. tumefaciens* genome (upper panel) and schematic of the 250bp region cloned into p*lacZ*290 to create a P*gcrA-lacZ* transcriptional reporter (into p*lacZ*290-P*gcrA* that can replicate into *E. coli* and *A. tumefaciens* cells). *lacZ* was represented with a dashed blue line as it is longer than *gcrA*. **(b)** p*lacZ*290-P*gcrA* was introduced into the indicated *A. tumefaciens* C58 derivatives. Cells were cultivated into ATGN+IPTG until cultures reached an $OD_{600}$~0.4. Cells were then washed and resuspended into ATGN+/-IPTG at an $OD_{600}$~0.05. Cultures were then incubated for 14 hours before samples were collected for β-galactosidase assays to evaluate the activity of the *gcrA* promoter in *A. tumefaciens*. Significant differences (ANOVA test coupled with Tuckey's multi-comparison test) are indicated by * (P<0.01) or ** (P<0.001). **(c)** p*lacZ*290-P*gcrA* was introduced into *E. coli* TOP10 cells carrying pBAD-RpoD$^{Cc}$-CcrM$^{Cc}$-WT (expressing the *C. crescentus* RpoD and CcrM proteins) or pBAD-RpoD$^{Cc}$-CcrM$^{Cc}$-mut (expressing the *C. crescentus* RpoD protein and an inactive variant of its CcrM protein). Cells were cultivated over-night into LB and then resuspended into LB+arabinose 0.3% at an $OD_{600}$~0.05. Cultures were then incubated for 3 hours before samples were collected for β-galactosidase assays to evaluate the activity of the *gcrA* promoter in *E. coli* cells expressing the *C. crescentus* RpoD$^{Cc}$ housekeeping Sigma factor together with an active or inactive CcrM$^{Cc}$ protein. Significant difference (Student's t-test) is indicated by * (P<0.0001). For all β-galactosidase assays (panels b and c), three biological replicates were done.

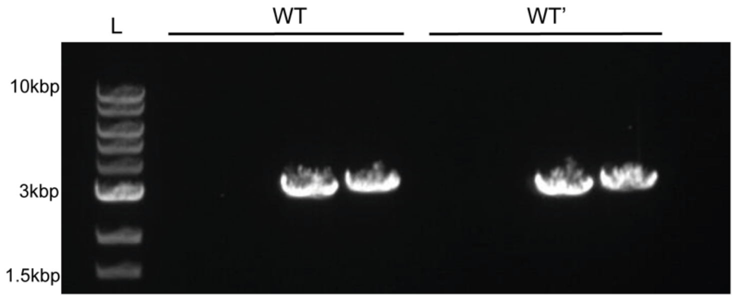


**Figure S13: PCR and gel electrophoresis analyses showing that JC2140 (WT) and JC2141 (WT') *A. tumefaciens* C58 cells display a unique dicentric chromosome as described in [4].** gDNA from WT and WT' cells were analyzed by PCR using F1/F2/R1/R2 primers as done before [4]. Detection of a PCR product with F1/R2 and F2/R1 primer pairs and no PCR product with F1/R1 and F2/R2 primer pairs indicates that JC2140 (WT) and JC2141 (WT') display a unique dicentric chromosome, which was also further confirmed by whole-genome sequencing and assembly. L: DNA ladder.

### SUPPLEMENTARY TABLES:

#### Table S1: Strains used in this study

| Strain name | Genotype | Description | Reference/Origin |
|---|---|---|---|
| ***Agrobacterium tumefaciens*** | | | |
| JC2140 (named WT in this article) | C58 | Wild-type *Agrobacterium tumefaciens* C58 with dicentric chromosome, pTiC58 and pAtC58. | [5] |
| JC2141 (named WT' in this article) | C58 *ΔtetRA*::a-*att*Tn*7* | Replacement of the *tetRA* locus with an artificial *att*Tn*7* site. Growth, motility and biofilm formation are identical to WT as shown in [5]. | [5] |
| JC2291 | WT' P*tac-ccrM* | | This study |
| JC2307 | WT' *ΔccrM* P*tac-ccrM* | | This study |
| JC2656 | WT' PT7-*ygfp-parB$^{MT1}$-parS$^{MT1}$* inserted between *Atu_0047* and *Atu_0048* | PT7-*ygfp-parB$^{pMT1}$-parS$^{pMT1}$* reporter near *Atu_0048 (*50kbp away from *ori1)* | This study |
| JC2657 | WT' PT7-*ygfp-parB$^{MT1}$-parS$^{MT1}$* inserted between *Atu_* 3973 and *Atu_3974* | PT7-*ygfp-parB$^{pMT1}$-parS$^{pMT1}$* reporter near *Atu_3973* (57kbp away from *ori2)* | This study |
| JC2777 | WT' PT7-*mcherry-parB$^{P1}$-parS$^{P1}$* inserted between *Atu_0047* and *Atu_0048* and PT7-*ygfp-parB$^{MT1}$-parS$^{MT1}$* inserted between *Atu_* 3973 and *Atu_3974* | PT7-*mcherry-parB$^{P1}$-parS$^{P1}$* reporter near *Atu_0048 (*50kbp away from *ori1)* and PT7-*ygfp-parB$^{pMT1}$-parS$^{pMT1}$* reporter near *Atu_3973* (57kbp away from *ori2)* | This study |
| JC2660 | WT' *ΔccrM* P*tac-ccrM* PT7-*ygfp-parB$^{MT1}$-parS$^{MT1}$* inserted between *Atu_0047* and *Atu_0048* | PT7-*ygfp-parB$^{pMT1}$-parS$^{pMT1}$* reporter near *Atu_0048 (*50kbp away from *ori1)* | This study |
| JC2661 | WT' *ΔccrM* P*tac-ccrM* PT7-*ygfp-parB$^{MT1}$-parS$^{MT1}$* inserted between *Atu_* 3973 and *Atu_3974* | PT7-*ygfp-parB$^{pMT1}$-parS$^{pMT1}$* reporter near *Atu_3973* (57kbp away from *ori2)* | This study |
| JC2836 | WT' *ΔccrM* P*tac-ccrM* PT7-*mcherry-parB$^{P1}$-parS$^{P1}$* inserted between *Atu_0047* and *Atu_0048* and PT7-*ygfp-parB$^{MT1}$-parS$^{MT1}$* inserted between *Atu_* 3973 and *Atu_3974* | PT7-*mcherry-parB$^{P1}$-parS$^{P1}$* reporter near *Atu_0048 (*50kbp away from *ori1)* and PT7-*ygfp-parB$^{pMT1}$-parS$^{pMT1}$* reporter near *Atu_3973* (57kbp away from *ori2)* | This study |
| ***Escherichia coli*** | | | |
| TOP10 | | Used for cloning procedures and β-galactosidase assays. | |
| JC1710 | S17.1 λpir | Used for plasmid transfers from *E. coli* to *A. tumefaciens* (conjugation). | [6] |
| ***Caulobacter crescentus*** | | | |
| NA1000 | Synchronizable CB15N wild-type strain | Used for PCR amplification of *ccrM$^{Cc}$* and *rpoD$^{Cc}$* | [7] |

**Table S2: Plasmids used in this study**

| Plasmid name | Description | Reference/origin |
|---|---|---|
| pTNS3 | Helper plasmid encoding the site-specific TnsABCD Tn7 transposition pathway ($Amp^R$) | [5] |
| pUC-miniTn7TGMPtac-HA | Mini-Tn7 vector containing *lacIq* and *tac* promoter ($Amp^R$ and $Gm^R$) | [5] |
| pUC-miniTn7TGMPtac-ccrM | *ccrM* ORF (*Atu_0794*) cloned into pUC-miniTn7TGMPtac-HA under the control of IPTG-inducible P*tac.* | This study |
| pNPTS138 | Suicide vector with *sacB* gene and *oriT* ($Km^R$) | D. Alley, unpublished |
| pNPTS138-ΔccrM | pNPTS138-derived suicide vector to delete the *ccrM* gene in the *A. tumefaciens* genome | This study |
| pWX963 | pNPTS138-derived suicide vector to integrate PT7-*ygfp*-$parB^{MT1}$-$parS^{MT1}$ near *Atu_0048* | [8] |
| pWX967 | pNPTS138-derived suicide vector to integrate PT7-*ygfp*-$parB^{MT1}$-$parS^{MT1}$ near *Atu_3973* | [8] |
| pWX995 | pNPTS138-derived suicide vector to integrate PT7-*mcherry*-$parB^{P1}$-$parS^{P1}$ near *Atu_0048* | [9] |
| pRXMCS-2 (also named pRX in figures) | Low copy number vector with RK2 origin and *oriT* ($Km^R$) | [10] |
| pRX-P*tac-ccrM* | P*tac-ccrM* from pUC-miniTn7TGMPtac-ccrM cloned into pRXMCS-2 | This study |
| p*lacZ*290 | Low copy number vector with RK2 origin and *oriT,* used to create *lacZ* transcriptional fusions ($Tet^R$) | [11] |
| p*lacZ*290-P*gcrA* | *gcrA* promoter region (250 bp upstream of *gcrA* ORF) cloned into p*lacZ*290 and controlling *lacZ* transcription | This study |
| pBAD24-CcrM-WT | Vector encoding the *Caulobacter crescentus* CcrM protein from an arabinose-inducible promoter ($Amp^R$) | [12] |
| pBAD24-CcrM-mut | Vector encoding a catalytically inactive *Caulobacter crescentus* CcrM(D31A) mutant protein from an arabinose-inducible promoter ($Amp^R$) | [12] |
| pBAD-$RpoD^{Cc}$-$CcrM^{Cc}$-WT | *Caulobacter crescentus rpoD* ORF (*CCNA_03142*) cloned into pBAD24-CcrM-WT and controlled by arabinose-inducible promoter | This study |
| pBAD-$RpoD^{Cc}$-$CcrM^{Cc}$-mut | *Caulobacter crescentus rpoD* ORF (*CCNA_03142*) cloned into pBAD24-CcrM-mut and controlled by arabinose-inducible promoter | This study |

**Table S3: Oligonucleotides used in this study**

| Primer name | Sequence (5'→3') | Used for: |
|---|---|---|
| ***Primers used for cloning procedures and strain constructions:*** | | |
| MS9 | GTACATATGGCAGCAGTTTTTCCGCTGG | Construction of pUC-miniTn7TGMPtac-ccrM |
| MS10 | CGAGATCTCTATTCAGCCTTTGCCATTTCAC | Construction of pUC-miniTn7TGMPtac-ccrM |
| Tet-Forward | ACATGTTGTATACCGGAAACTGATTGCAC | Verification of P*tac-ccrM* insertion into genome of JC2291 |
| Tn7R109 | CAGCATAACTGGACTGATTTCAG | Verification of P*tac-ccrM* insertion into genome of JC2291 |
| MS1 | GCCAAGCTTCAATTTTGCTTCGAAGGTGGCGT | Constructions of pNPTS138-ΔccrM and strain JC2307 |
| MS2 | TGCGGATCCTGGTACTCGCTCCATACGCTTAC | Construction of pNPTS138-ΔccrM |
| MS3 | GCTGGATCCTCGAGAGCCGTAGAGGCTGGAG | Construction of pNPTS138-ΔccrM |
| MS4 | GGGGCTAGCTACCGCTCCAGTCCTGCGCAAT | Constructions of pNPTS138-ΔccrM and strain JC2307 |
| MS17 | TCCTTCACCTCGAAGGGCAC | Construction of strain JC2307 |
| MS40 | CCAGAATTCGCTGTGGTATGGCTGTGCAG | Construction of pRX-P*tac-ccrM* |
| MS41 | GGCAGATCTTAGCTAGCTCATTAAGCGTAATCTGGAAC | Construction of pRX-P*tac-ccrM* |
| MS46b | CCACTGCAGGCCGCTTCTCCGCTTTTC | Construction of p*lacZ*290-P*gcrA* |
| MS47 | TGGGAATTCCTTTCGAGGCCCAAGGTG | Construction of p*lacZ*290-P*gcrA* |
| LC26 | GCGCAAGCTTAGGAGGAATTCACCATGAGCAACAATTCCTCGGCCGA | Construction of pBAD-RpoD$^{Cc}$-CcrM$^{Cc}$-WT and pBAD-RpoD$^{Cc}$-CcrM$^{Cc}$-mut |
| LC29 | AAAAAAAAGCTTTTACGAGTCCAGGAAGCTGCGCA | Construction of pBAD-RpoD$^{Cc}$-CcrM$^{Cc}$-WT and pBAD-RpoD$^{Cc}$-CcrM$^{Cc}$-mut |
| LC32 | GCGAGGACCTCTCGGACGCCG | Construction of pBAD-RpoD$^{Cc}$-CcrM$^{Cc}$-WT and pBAD-RpoD$^{Cc}$-CcrM$^{Cc}$-mut |
| LC33 | GCGGATGTCGGTGACCGACTGG | Construction of pBAD-RpoD$^{Cc}$-CcrM$^{Cc}$-WT |

| | | |
|---|---|---|
| | | and pBAD-RpoD$^{Cc}$-CcrM$^{Cc}$-mut |
| ***Primers used for qRT-PCR:*** | | |
| MS90 | CGCTTCGTGCAGAAGGACT | *hemF* |
| MS91 | GCACGCCGACCTTTTCGAA | |
| MS80 | GCGCTCTGATGCGGGTTTT | *purH* |
| MS81 | AACTGGTTTGCCGAAGCACTG | |
| MS84 | CTGCCGTCCTTGCCGGAAA | *yidC* |
| MS85 | AGCCGGTAATTCCCCCTGTATT | |
| MS154 | GGACTGGCTGTTCCCGATTT | *ccrM* |
| MS155 | CTGTGTGGGATGCACCTTCTT | |
| MS100 | GCCGACGCTCGTCGTCGTAA | *repA*$^{Ch2}$ |
| MS101 | CGATCCCCAGGCCAGTCT | |
| MS102 | GGGAACCGCATCTGCAGTTT | *repB*$^{Ch2}$ |
| MS103 | GTCGCCCAGGGTCAGGAA | |
| MS104 | GCGGGCATAACGTTTTCCGTT | *repC*$^{Ch2}$ |
| MS105 | GGCATGGCCGAACAGACATT | |
| MS98c | AGTTCCGTCAGCGTCAGG | *gcrA* |
| MS99b | GCGTCGAGACTGCAGAAGTTC | |
| MS146 | GGAGCGTGGTAGACCGGT | *ftsZ*$^{AT}$ |
| MS147 | CAGCAGCCTTCTCAGGACTT | |
| ***Primers used to test if the C58 derivatives (WT/WT') are fusion strains with a dicentric chromosome*** | | |
| F1 | TGTCTGGGTTCTGGAATTCGACGC | [4] |
| R1 | AGGTTCCGTGGTATAGTTGTAGGC | [4] |
| F2 | CTTGATCCAGAGTGATTTCGACGC | [4] |
| R2 | CCTTGTGAACAACGCCTTTGACCC | [4] |

**Table S4 (Excel Supplementary file): RNA-Seq results comparing the transcriptome of JC2307 (*ΔccrM* P*tac-ccrM*) cells cultivated in ATGN +/- IPTG (7 hours) and of JC2140 (WT) cells cultivated in ATGN-IPTG.** Replicon NC_003062.2 corresponds to Ch1 in non-fusion strains, replicon NC_003063.2 corresponds to Ch2 in non-fusion strains, replicon NC_003064.2 corresponds to pAt and replicon NC_003065.2 corresponds to pTi. Activated (FC>2) genes are highlighted in green. Repressed (FC>2) genes are highlighted in red. Adjusted P-values >0.01 (non-significant changes) are highlighted in grey. Genes of particular interest that are discussed in this study are highlighted in yellow. For COG annotations, the COG table in json format for *Agrobacterium tumefaciens* (Agrobacterium_fabrum_C58) was downloaded from the NCBI Database of Clusters of Orthologous Genes ( https://www.ncbi.nlm.nih.gov/research/cog/) [13]. The COG json table was reformatted into text format and gene identifiers were matched to COG terms via a customised Perl script. For missing terms, the NIH GenBank database of *Agrobacterium tumefaciens* protein sequences was used [14]. GANTC motifs in 200 bp sequences upstream of the annotated start codon of each gene were searched using an ad hoc Perl script to find motif matches in FASTA-formatted sequences.

**Table S5 (Excel Supplementary file): Lists of genes that were significantly mis-regulated (FC>2 and adjusted P-value<0.01) when comparing the transcriptome of JC2307 (*ΔccrM* P*tac-ccrM*) cells cultivated in ATGN +/- IPTG (7 hours) and genes that can be considered as part of the "direct regulon" of CcrM (FC>2, adjusted P-value<0.01 and minimum one GANTC motif in the 200 bp upstream of ORF).** Activated genes are highlighted in green. Repressed genes are highlighted in red. Genes of particular interest that are discussed in this study are highlighted in yellow. In the "direct regulon" tab, orthologous genes that also belong to the "direct regulon" of the *Caulobacter crescentus* NA1000 CcrM protein (from [15]) are highlighted in blue and orthologous genes that also belong to the "direct regulon" of the *Brevundimonas subvibrioides* CcrM protein (from [16]) are highlighted in orange.

To search for genes orthologous to genes that belong to the "direct regulon" of CcrM in *A. tumefaciens* in the *C. crescentus NA1000* (NC_011916.1) or *B. subvibrioides* (NC_014375.1) genomes, the *blastP* alignment program [17] was used. In short, blast-formatted sequence databases were created for the three genomes and a blastP search was performed in command-line mode, with the following parameter options: "-num_alignments 5 -subject_besthit -outfmt "6 qacc qlen qstart qend sacc slen sstart send length pident evalue bitscore score mismatch gapopen gaps stitle". For further validation and complementary information, *blastP* searches were supplemented with searches based on COG identifiers between genes in the different species. This analysis revealed that only 4 genes belonging to the *C. crescentus* "direct" CcrM regulon also belonged to the *A. tumefaciens* "direct" CcrM regulon (1$^{st}$ blatsp matches and COG clusters). Similarly, only two genes belonging to the *B. subvibrioides* "direct" CcrM regulon also belonged to the *A. tumefaciens* "direct" CcrM regulon (2$^{nd}$/3$^{rd}$ best blastp matches).

| Gene ID | Gene Name | Gene Type | Replicon | Gene Start | Gene End | Strand |
|---|---|---|---|---|---|---|
| *Atu8080-1* | repE-linear-1 | ncRNA | NC_003063.2 | 1024262 | 1024335 | + |
| *Atu8080-2* | repE-linear-2 | ncRNA | NC_003063.2 | 1024298 | 1024375 | + |
| *Atu8080-3* | repE-linear-3 = $RepE^{Ch2}$ | ncRNA | NC_003063.2 | 1024387 | 1024464 | + |
| *Atumisc_RNA_3-pAt* | repE-pAt-3 | ncRNA | NC_003064.2 | 2398 | 2478 | - |
| *Atumisc_RNA_2-pAt* | repE-pAt-2 | ncRNA | NC_003064.2 | 2595 | 2675 | - |
| *Atumisc_RNA_1-pAt* | repE-pAt-1 | ncRNA | NC_003064.2 | 2542 | 2605 | - |
| *Atumisc_RNA_1_pTi* | repE-pTi | ncRNA | NC_003065.3 | 54724 | 54801 | - |

**Table S6: Custom *Atu* small RNA genes added into the annotation during RNA-Seq analyses.** The presence of these transcripts was detected in the sequenced RNA during the RNASeq analyses using RNA from strains JC2140 and JC2307 as described in Fig.S7.

### SUPPLEMENTARY METHODS:

#### Plasmid constructions

Construction of **p*lacZ*290-P*gcrA***: The 250 bp promoter region upstream of the *gcrA* (*Atu0426*) ORF of the *A. tumefaciens* C58 strain (JC2140) was amplified using primers MS46b and MS47, digested with PstI/EcoRI and ligated into PstI/EcoRI-digested p*lacZ*290.

Constructions of **pBAD-RpoD$^{Cc}$-CcrM$^{Cc}$-WT** and **pBAD-RpoD$^{Cc}$-CcrM$^{Cc}$-mut**: The *rpoD$^{Cc}$* gene (*CCNA_03142)* was amplified from *C. crescentus* NA1000 gDNA using primers LC26 and LC29, creating a 5'AGGAGG3' *E. coli* ribosome binding site in frame with *rpoD$^{Cc}$*. The PCR product and the pBAD-RpoD$^{Cc}$-CcrM$^{Cc}$-WT or pBAD-RpoD$^{Cc}$-CcrM$^{Cc}$-mut vectors were then digested with the HindIII enzyme and ligated together. The insert sequence and orientation were checked using primers LC32 and LC33.

#### β-galactosidase assays

β-galactosidase assays were adapted from [18]. Briefly, the $OD_{600nm}$ of each culture was measured and a given volume (V, in mL and below 500μL) of each culture was collected. Z-buffer (without β-mercaptoethanol) was then added to each culture sample to reach a final volume of 1mL. 30μL of 0.05% SDS and 60μL chloroform were then added into these samples, which were then mixed vigorously. Samples were then incubated at 28°C for 5 minutes to reach optimum temperature. The β-galactosidase reaction was then started when 200μL of ONPG at 4mg/mL was added. After a given reaction time (t, in minutes), the reaction was stopped with the addition of 500μL of 1M $Na_2CO_3$ and samples were briefly centrifuged before measuring the $A_{420nm}$. Miller Units (MU) were calculated using the following formula: MU = $1000*A_{420}/(OD_{600}*t*V)$. 3 biological replicates were used for each strain/condition.
